# PGC-1α overexpression partially rescues impaired oxidative and contractile pathophysiology following volumetric muscle loss injury

**DOI:** 10.1101/535328

**Authors:** William M. Southern, Anna S. Nichenko, Kayvan F. Tehrani, Melissa J. McGranahan, Laxminarayanan Krishnan, Anita E. Qualls, Nathan T. Jenkins, Luke J. Mortensen, Hang Yin, Amelia Yin, Robert E. Guldberg, Sarah M. Greising, Jarrod A. Call

**Affiliations:** Department of Kinesiology, University of Georgia, Athens, GA 30602, USA; Regenerative Bioscience Center, University of Georgia, Athens, GA 30602, USA; Parker H. Petit Institute for Bioengineering & Bioscience, Georgia Institute of Technology, Atlanta, GA 30332, USA; Center for Molecular Medicine, University of Georgia Athens, GA 30602, USA; Department of Biochemistry and Molecular Biology, University of Georgia, Athens, GA 30602, USA; Knight Campus for Accelerating Scientific Impact, University of Oregon, Eugene, OR 97403, USA; School of Kinesiology, University of Minnesota, Minneapolis, MN, 55455, USA

**Keywords:** Mitochondrial Biogenesis, Vasculature, Wheel Running, Muscle Strength, Muscle Function, Neuromusculoskeletal Injury, Orthopaedic Trauma, Skeletal Muscle Injury

## Abstract

Volumetric muscle loss (VML) injury is characterized by a non-recoverable loss of muscle fibers due to ablative surgery or severe orthopaedic trauma, that results in chronic functional impairments of the soft tissue. Currently, the effects of VML on the oxidative capacity and adaptability of the remaining injured muscle are unclear. A better understanding of this pathophysiology could significantly shape how VML-injured patients and clinicians approach regenerative medicine and rehabilitation following injury. Herein, the data indicated that VML-injured muscle has diminished mitochondrial content and function (i.e. oxidative capacity), loss of mitochondrial network organization, and attenuated oxidative adaptations to exercise. However, forced PGC-1α over-expression rescued the deficits in oxidative capacity and muscle strength. This implicates physiological activation of PGC1-α as a limiting factor in VML-injured muscle adaptive capacity and provides a mechanistic target for regenerative rehabilitation approaches to address the skeletal muscle dysfunction.

---

Oxidative capacity is a cornerstone of skeletal muscle health, and for the past 40 years, we have known that the most robust physiologic adaptation to regularly scheduled physical activity (i.e., exercise/overload training) is an increase in oxidative capacity(1, 2). Improvements in muscle oxidative capacity are made possible with exercise training through adaptations affecting the density and function of the intramuscular mitochondrial network. The signaling pathways that initiate and coordinate mitochondrial improvements with exercise are complex, but advancements in molecular biology in the last two decades have revealed many of the key players involved (see for review(3)). Most notably, the transcription factor PGC-1α (peroxisome proliferator-activated receptor gamma, coactivator 1 alpha) is considered a critical molecular modulator of skeletal muscle oxidative plasticity because it regulates gene expression patterns for expansion of the mitochondrial network (i.e. mitochondrial biogenesis), angiogenesis, and motor neuron associated adaptations with exercise training(4–6). Expansion of the vasculature and mitochondrial network with exercise training enhances the functional capacity of the muscle (e.g., fatigue resistance), and in general, this physiologic type of acclimation is considered beneficial for human performance and health(7, 8).

Large-scale skeletal muscle trauma, such as volumetric muscle loss (VML) injury, is unique in that the muscle is not able to regenerate muscle fibers with endogenous repair systems and as a result, cannot fully recover strength. The loss of muscle function (i.e., contractility) can exceed the loss of tissue mass(9), and this permanent functional deficit leaves patients with lifelong disability(10) for which there is currently no corrective physical rehabilitation guidelines. Furthermore, the extent to which the remaining skeletal muscle can adapt to rehabilitation is unclear. Recent preclinical work has investigated various models of ‘physical rehabilitation’ following VML in the forms of voluntary wheel running(11–14), forced treadmill running(14), chronic-intermittent electrical nerve stimulation and/or passive range of motion exercises(15) and collectively found modest contractile adaptations are possible. However, clinical reports have indicated that patients only see moderate improvements and then hit the ceiling where further physical therapy, no matter the type or intensity, does not result in an increase in function(16–18). Collectively, investigations of physical rehabilitation following VML have resulted in, at best, modest improvements in muscle function following VML injury without any physiological rationale or mechanistic understanding for the lack of significant response.

Overwhelmingly, investigations of preclinical outcome measures have focused on histologic and/or contractile aspects of the injured muscle(9) with very few investigations focusing on any aspect of the oxidative plasticity of the remaining muscle. Noteably, Aurora and colleagues reported a reduced metabolic gene response (i.e., PGC-1α and SIRT-1) in VML-injured muscle following voluntary wheel running compared to uninjured muscle(11). Furthermore, Greising and colleagues showed that mitochondrial respiration rates measured from both the VML-injured and contralateral uninjured mouse muscles were less than that of injury naïve mice suggesting a more chronic and systemic effect of VML injury on muscle oxidative capacity(15). Therefore, VML-injured patients may be susceptible to lifelong impairments in skeletal muscle oxidative capacity that can result in additional comorbidities as systemic reductions in skeletal muscle oxidative capacity are associated with an increased risk for a host of disorders such as diabetes(19) and cardiovascular disease(20).

One apparent reason for the lack of investigation into oxidative capacity of VML-injured muscle is the difficulty in assessing physiological mechanisms of mitochondrial function. Herein, we have combined high-resolution mitochondrial respirometry with mitochondrial enzyme kinetics and 2-photon microscopy to overcome this difficulty. The seminal work by Glancy et al.(21, 22) utilized 2-photon imaging to characterize the highly organized mitochondrial network and brought to light the importance of the network as it relates to the function of the mitochondria. An important innovation of our work was to use the 2-photon microscopy to investigate changes in the structural integrity of the mitochondrial network as this aspect of mitochondrial physiology is likely disrupted following VML injury. The combination of all of these proven techniques has allowed us to extensively characterize multiple aspects of mitochondrial physiology in VML-injured muscle, which provides a new and unique prospective of the impacts of VML on mitochondria.

Repair of VML injury has been approached extensively by biomedical engineering and regenerative medicine experts over the past decades(23), yet no clear indication of clinically meaningful therapies capable of significantly improving skeletal muscle function have emerged. We posit that a dysfunctional oxidative capacity contributes to a poor regenerative niche within the VML-injured muscle that ultimately limits regenerative approaches. Here we report that the pathophysiology of the remaining muscle after VML includes widespread impairments in mitochondrial structure and respiratory function, and an insensitivity to physiological stimuli known to enhance mitochondrial structure and function (i.e., exercise). We identify the activation of the transcription factor PGC-1α as the limiting factor to exercise-induced adaptation and demonstrate the forced over-expression of PGC-1α rescue a substantial portion of the VML pathophysiology.

## Results

### Muscle Function and Oxidative Deficits following Multi-muscle Model of VML Injury

With the goal of developing treatment strategies capable of improving the long-term functional deficits in VML injured muscle tissue, we sought to elucidate the early effects of VML on contractility and oxidative capacity of the remaining skeletal muscle. In order to best characterize the impact of VML on skeletal muscle, we used a model of VML injury that has been shown to reproducibly recapitulate the injury on primary hind limb locomotor muscles (i.e., the plantar flexor muscles [gastrocnemius, soleus, plantaris]) and results in a chronic functional deficit(15). In agreement with prior reports,(24–26) muscle contractile function was reduced by ~80% when compared to uninjured muscles for injury naïve mice at both 3 and 7 days post-injury, even after normalizing to the injured muscle mass (P<0.001; Fig. 1a-b, Table 1).

**Figure 1:**
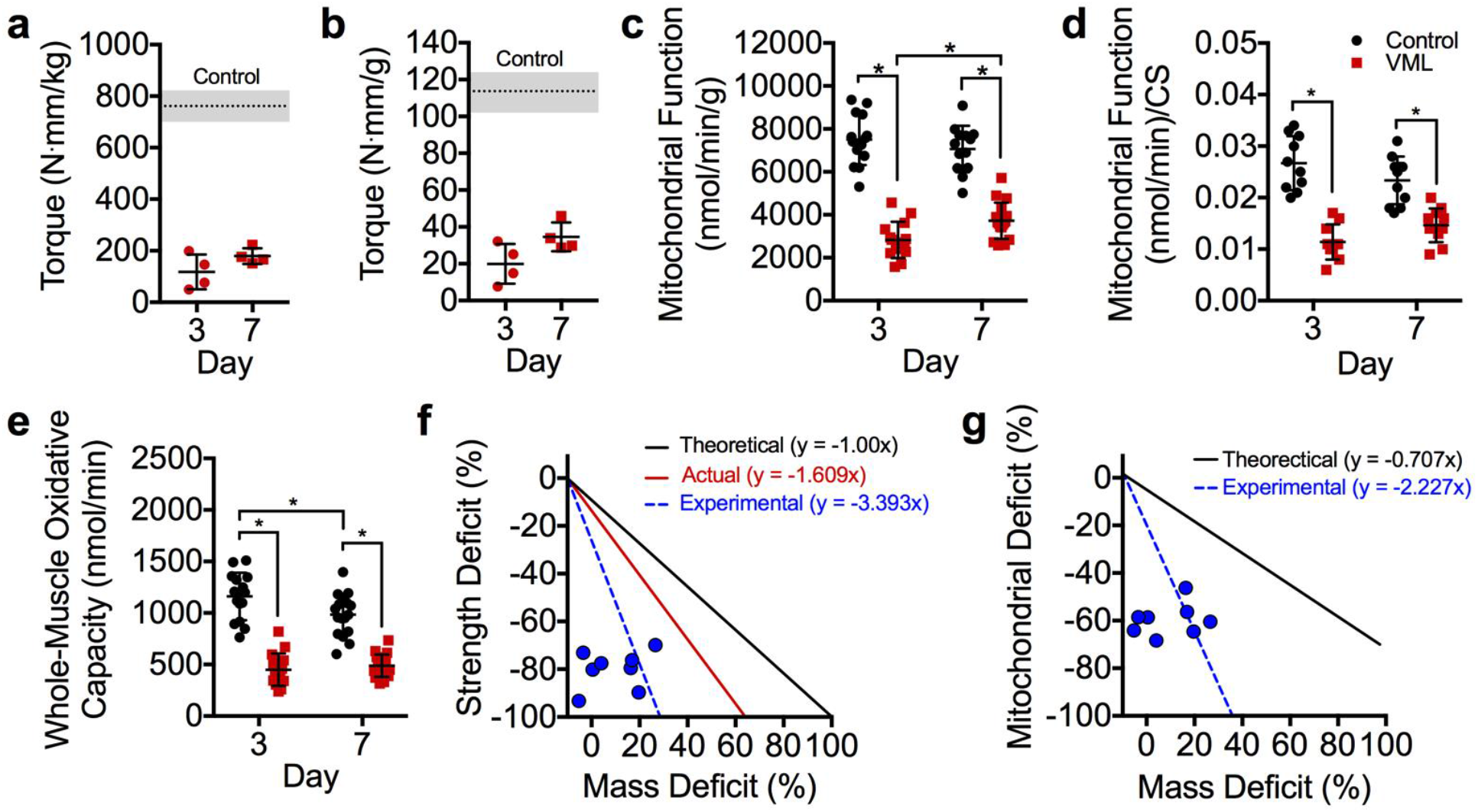
Early, 3 and 7 days, following VML injury the muscle remaining undergoes remarkable alterations in both the contractile and oxidative components of the muscle. Peak isometric torque normalized to (**a**) body mass and (**b**) plantarflexor mass is significantly reduced at both 3 and 7 days post-VML compared to control (dotted line, SD indicated by shading) (different from control in both a and b, P<0.001). (**c**) Mitochondrial function normalized by grams wet weight of permeabilized muscle fibers (n≥15 permeabilized fiber bundles from n=4 mice for each group) is also significantly reduced following VML injury. (**d**) Mitochondrial respiratory function normalized to citrate synthase enzyme activity. (**e**) Extrapolation of mitochondrial respiration rates (panel c) to entire muscle mass (see Table 1). (**f**) Theoretical 1:1 relationship between muscle mass and contractile function (black); compolation of previously published work (see for review(9)) showing the relationship between the volume of muscle tissue removed at injury and the contractile function 1-4 months after VML (red); experimental data collected herein showing the relationship between muscle mass lost 3-7 days after VML and contractile function (blue, hashed lines and dots). (**g**) Theoretical 0.82:1 relationship between deficits in muscle mass and whole muscle oxidative capacity determined from various degenerative etiology (i.e., denervation, aging, cachexia, immobilization, heart failure, ischemia reperfusion injury, and critical illness) (black); the experimental data collected herein between deficits in muscle mass and whole muscle oxidative capacity determined herein (blue, hashed lines and dots, see panel **e**). Data analyzed by one- or two-way ANOVA, *P<0.05. Throughout, error bars represent means ± SD.

**Table 1.** Mouse gastrocnemius muscle masses and citrate synthase activity

| **Acute Study** | VML Day 3 |  | VML Day 7 |  |
| --- | --- | --- | --- | --- |
|  | Contralateral | Injured | Contralateral | Injured |
| Gastrocnemius Mass (mg) | 160.3 ± 17.1 | 155.1 ± 20.1 | 155.3 ± 9.9 | 129.8 ± 6.3<sup>a</sup> |
| Citrate Synthase (μmol/min/g) | 569.9 ± 24.7 | 529.3 ± 29.7<sup>b</sup> | 563.0 ± 16.5 | 539.0 ± 19.0<sup>b</sup> |

| **Wheel Running Study** | Control |  | VML |  | VML |  |
| --- | --- | --- | --- | --- | --- | --- |
|  | No Run | Run | Contralateral | Injured | Contralateral + Run | Injured + Run |
| Gastrocnemius Mass (mg) | 174.9 ± 13.7 | 174.8 ± 7.4 | 162.3 ± 10.5 | 129.7 ± 5.9<sup>c</sup> | 157.3 ± 12.5 | 130.0 ± 23.1<sup>c</sup> |
| Citrate Synthase (μmol/min/g) | 557.8 ± 29.1 | 536.5 ± 91.9 | 487.6 ± 100.8 | 479.9 ± 50.2<sup>e</sup> | 535.0 ± 58.6 | 622.8 ± 103.3<sup>d</sup> |

| **PGC-1α Transfection Study** | Control |  | VML |  | VML |
| --- | --- | --- | --- | --- | --- |
|  | No PGC-1α | PGC-1α | Contralateral | Injured | Injured + PGC-1α |
| Gastrocnemius Mass (mg) | 174.9 ± 13.7 | 176.8 ± 14.7 | 170.6 ± 14.0 | 90.8 ± 12.0<sup>f</sup> | 104.4 ± 17.6<sup>f</sup> |
| Citrate Synthase (μmol/min/g) | 557.8 ± 29.1 | 554.2 ± 85.8 | 657.8 ± 68.5<sup>g</sup> | 512.4 ± 98.9 | 621.5 ± 89.1<sup>h</sup> |

Values are means ± SD; Data analyzed by one- or two-way ANOVA within each study
<sup>a</sup>Different from all other groups
<sup>b</sup>Main effect of injury: Contralateral > Injured
<sup>c</sup>Different from all uninjured limbs
<sup>d</sup>Different from control run, VML contralateral, VML injured, VML contralateral + Run
<sup>e</sup>Different from control no run
<sup>f</sup>Different from all uninjured limbs
<sup>g</sup>Different from control no PGC-1α, control PGC-1α, and VML injured
<sup>h</sup>Different from VML injured

To determine if VML affects muscle oxidative capacity early after injury, mitochondrial function was assessed at 3 and 7 days post-VML via high-resolution respirometry of permeabilized fibers isolated from portions of the remaining muscle adjacent to the injury site. Mitochondrial function of the injured muscle was significantly reduced by ~60% and ~50% compared to the uninjured contralateral limb at 3 and 7 days post-injury, respectively (Fig. 1c-e). Mitochondrial function was still lower than uninjured muscle even after accounting for concomitant reductions in mitochondrial content (i.e., citrate synthase; Fig. 1d, Table 1). Together, these data indicate the pathophysiology of VML-injured muscle includes contractile and metabolic dysfunction independent of muscle mass or mitochondrial content.

A disproportionate loss in contractile function for the volume of muscle removed during a VML injury has been previously documented (see for review(9)), and is likely caused by the extensive disruptions in muscle architecture, vasculature, and motor neurons caused by the injury. Our results are in agreement with this position (Fig. 1F); however, a similar analysis between mitochondrial function and volume of tissue removed has not been conducted, and results from such an analysis could bolster support for investigating aspects of mitochondrial structural integrity as a contributor to metabolic dysfunction in VML-injured muscle. We conducted an analysis from previously published non-VML conditions where muscle volume decreases due to injury or disease [i.e., denervation(27–31), aging(32–35), cachexia(36–38), immobilization(39), heart failure(40), ischemia reperfusion injury(41), and critical illness(42)]. Based on these reports, the deficit in oxidative capacity was similar to the deficit in muscle volume (ratio: 0.82:1). In contrast, we find a very large oxidative deficit in the VML injured muscle compared to the amount of muscle lost from the injury (Fig. 1g), suggesting that oxidative capacity disproportionately decreases in relation to the deficit in muscle tissue following VML.

### Substantial Structural Disruptions in the Mitochondrial Network in the Muscle Remaining after VML Injury

One of the many important determinants of mitochondrial function within muscle is the location and structural organization of the mitochondrial network(21, 22, 43). We posited that severe mechanical damage to muscle from a VML injury, likely leads to structural disruptions in the remaining mitochondrial network, which contributes to mitochondrial dysfunction. We utilized a transgenic mouse model with ubiquitously expressed mitochondrial Dendra2 green monomeric fluorescent protein ((44), Jackson Laboratory, #018385) and quantitative 2-photon microscopy to evaluate the mitochondrial network in the remaining muscle after VML injury to the tibialis anterior (TA) muscle. The TA muscle was utilized here to overcome the uneven imaging plane and multi-pennate nature of the gastrocnemius muscle. 3D reconstructions of the mitochondrial network were generated for uninjured muscle and injured muscle at 3, 7, and 28 days post-injury (Fig. 2). The mitochondrial network was substantially disorganized, specifically at 3 and 7 days after the injury compared to uninjured muscle (Fig. 2b-c).

**Figure 2:**
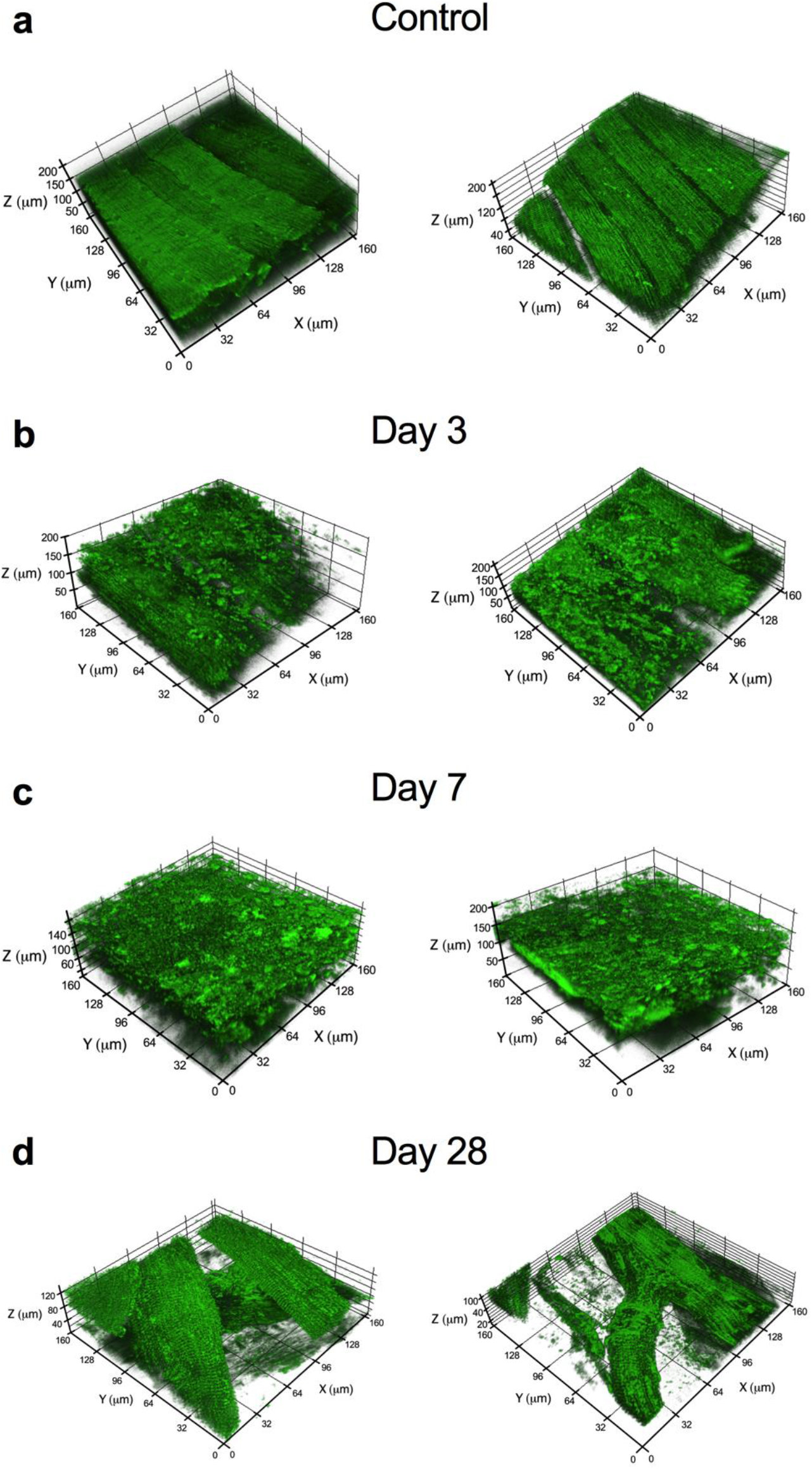
Significant changes and structural disruptions in the muscle remaining following VML injury. The effects of VML on mitochondrial organization were evaluated at various time points in control muscle and muscle remaining after VML injury. Two biological replicates representative 3D reconstructions of the mitochondrial network as observed in uninjured control muscle (**a**) and from the injury boundary in VML injured muscle at (**b**) 3, (**c**) 7, and (**d**) 28 days post-VML. Longitudinal striations are visible in uninjured control muscle indicating the high level of mitochondrial network organization, which is absent in the VML-injured muscle at 3 and 7 days after the injury.

Next, we quantified the mitochondrial network organization within each 3D mitochondrial network reconstruction. The mitochondrial network in uninjured muscle qualitatively appeared highly organized, with mitochondrial structures linearly aligned to two primary axes: parallel and perpendicular to muscle fiber orientation (Fig. 3a-b). Using this aspect of mitochondrial networks, an angular Fourier filtering (AFF) analysis method was developed to quantify mitochondrial structural organization by assessing the strength of structural alignment to a given angle. Our filtering method scans in a 360 degree arc and quantifies the strength of alignment of mitochondrial structures to various angles within the arc. With this analysis, well-connected and well-defined linear mitochondrial structures will have strong alignment to two specific angles/axes that are parallel and perpendicular to the longitudinal axis of the fiber. To quantify the organization of mitochondrial structures, the ratio of peak and average alignment strength was then calculated across each 3D mitochondrial network reconstruction (Fig. 3c). Mitochondrial network organization was reduced by ~50% and 70% compared to uninjured muscle at 3 and 7 days post-injury, respectively (P<0.001; Fig. 3c-d), in line with reductions in mitochondrial function, yet more severe than reductions in mitochondrial content. Even at 28 days post-injury, the mitochondrial network organization was still ~45% lower than uninjured muscle, which correlates with VML-induced chronic disruptions in oxidative capacity within the remaining muscle, independent of mitochondrial content (Fig. 2d & 3d). We suspect the disruptions in mitochondrial structure following VML injury are consistent with the observed deficits in mitochondrial function in the remaining muscle.

**Figure 3:**
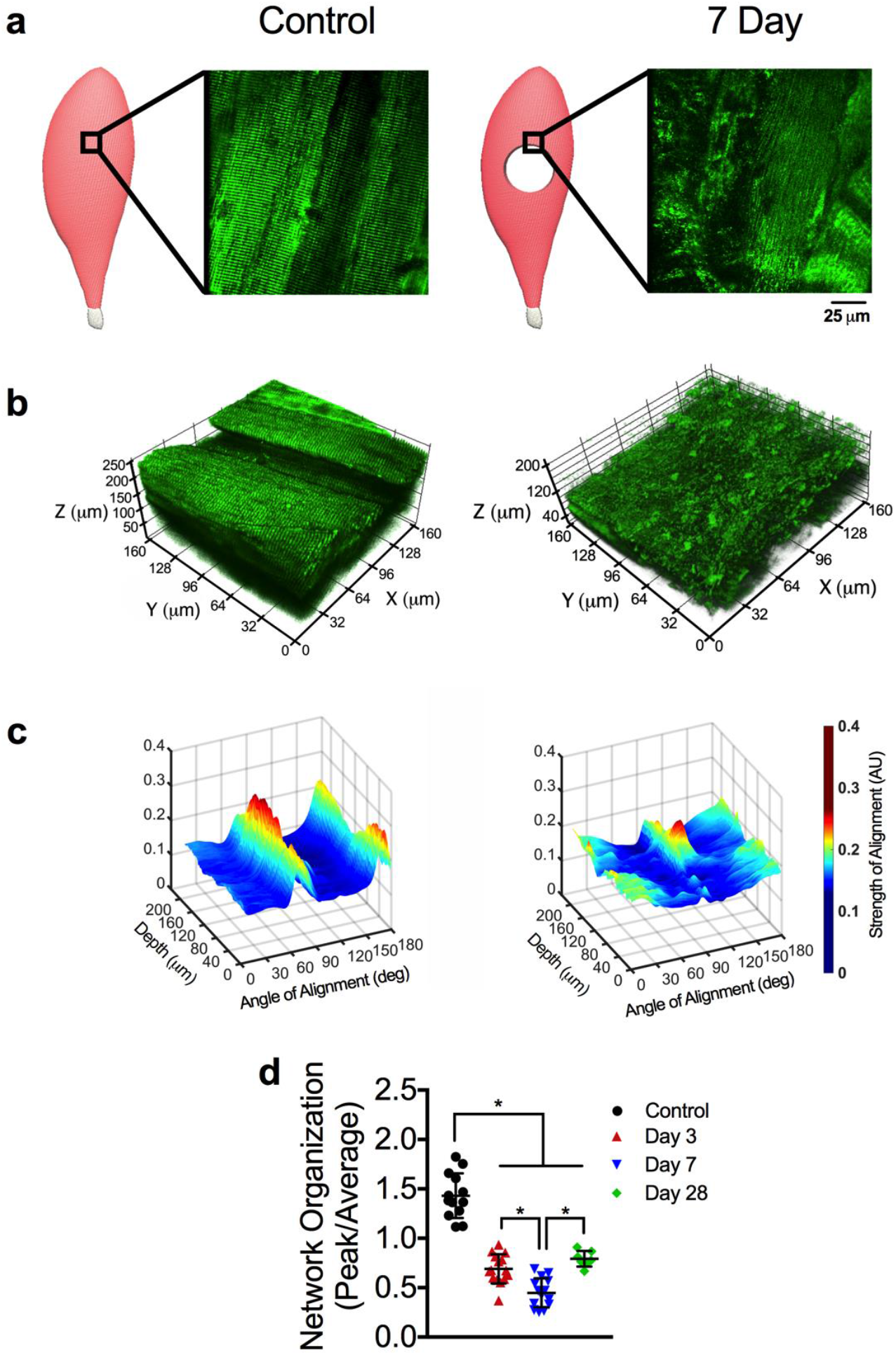
The mitochondrial network organization is altered by VML injury. (**a**) Schematics representing control and VML injured TA muscle, showing the region of interest evaluated during imaging and representative 2D images from the respective regions from transgenic mice that ubiquitously expressed mitochondrial Dendra2 GFP. (**b**) Representative 3D reconstruction of mitochondrial network adjacent to the VML injury site from control and VML injured TA muscles. (**c**) Representative 3D surface plots showing angle and alignment strength of the 3D mitochondrial network depicted. (**d**) Quantification of mitochondrial network organization (peak alignment/average alignment) at various time points after injury (n>10 z-stacks per muscle, per group). Data analyzed by one-way ANOVA, *P<0.05. Error bars represent means ± SD.

To assess the impact of VML on the mitochondrial structure across the entire muscle, we imaged a VML-injured muscle 7 days after injury at various distances (i.e., 0, 0.5, 1.5 2.0 and 2.5 mm) from the border of the injury in the proximal direction toward the origin of the muscle (Fig. 4a-g). Compared to uninjured control muscle, the mitochondrial network organization in the VML-injured muscle was reduced by ~40-50% at 0-2 mm away from the injury border, respectively (Fig. 4h). However, at 2.5 mm from the injury site (Fig. 4g), the mitochondrial organization returned to more than 90% of control, suggesting this was the outer edge of the mitochondrial damage from the VML. These findings suggest that the confounding impact of the VML injury on mitochondrial organization and function was observed beyond the border (i.e. 2mm) of the initial defect area, which potentially could result in diminished oxidative capacity in a significant portion of the muscle.

**Figure 4:**
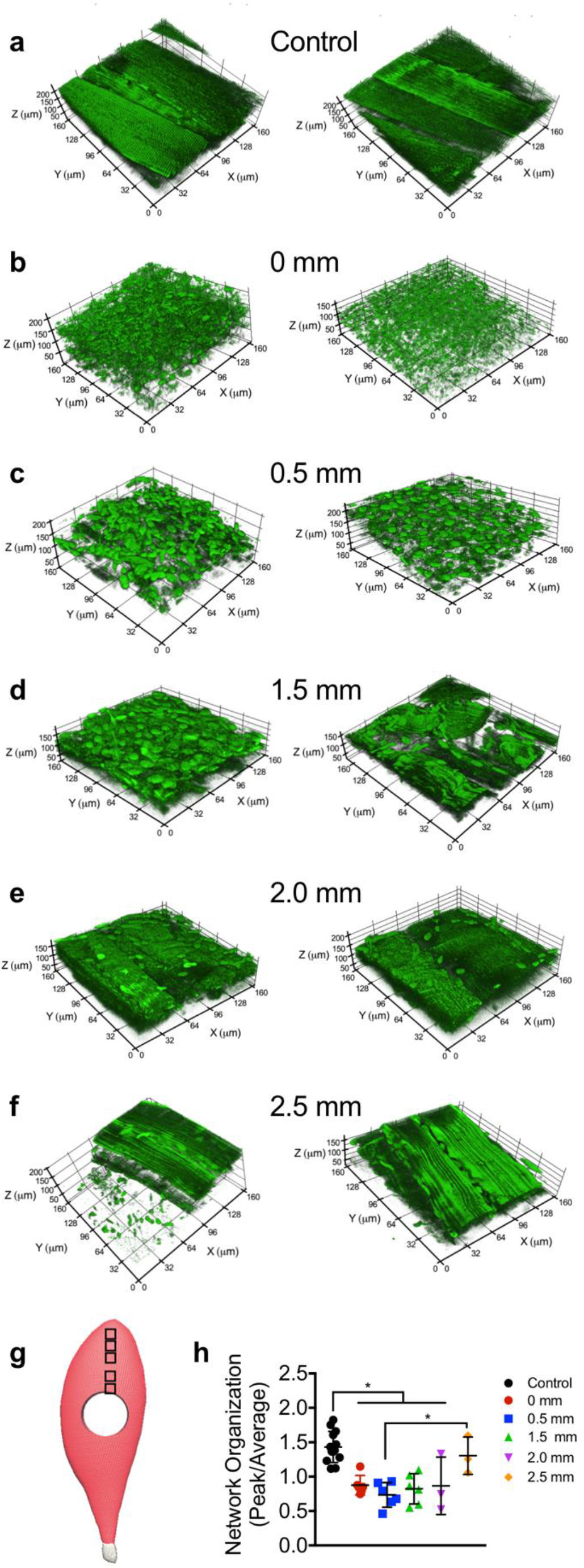
Structural alterations in mitochondrial network extend well beyond the site of VML injury. The effects of VML on mitochondrial organization in control and muscle remaining 7 days after VML injury were evaluated at various distances from the injury site, two representative 3D reconstructions of mitochondrial networks are presented for each experimental condition. Representative images from (**a**) uninjured control and (**b**) 0 mm, (**c**) 0.5 mm, (**d**) 1.5 mm, (**e**) 2.0 mm, and (**f**) 2.5 mm away from proximal VML injury border. (**g**) Schematic showing an injured TA with boxes to indicating the imaging sites at increasing distances from the border of the VML injury toward the origin of the muscle. (**h**) Quantification of mitochondrial network organization (peak alignment/average alignment) at various distances from VML injury site (control n=10, 0 mm; 0.5 mm, 1.5 mm n=6; 2.0 mm, 2.5 mm n=3 z-stacks per muscle). Data analyzed by one-way ANOVA, *P<0.05. Error bars represent means ± SD.

### VML-injured Muscles have Impaired Oxidative but not Contractile Plasticity

Exercise training is the most robust physiological stimulus to improve oxidative capacity within skeletal muscle. To facilitate improvements in oxidative capacity of the injured muscle, early exercise-based rehabilitation was implemented in the form of voluntary wheel running, which began 72 hours after the injury and lasted for 4 weeks. Uninjured mice demonstrated expected improvements in mitochondrial function indicating that running was sufficient to produce oxidative adaptations (Fig. 5a-b, Table 1). However, despite the fact that the same functional load was placed on the muscle (i.e., similar distance run; Fig. 5c), there were no oxidative adaptations in the remaining muscle following VML injury, suggesting that the mitochondria in the remaining muscle are not plastic (Fig. 5a & d). Muscle contractions and subsequent intercellular perturbations in ATP homeostasis that occur from muscle loading during exercise is a critical signal for mitochondrial adaptations to exercise training. A possible explanation for the lack of mitochondrial adaptations to wheel running is reduced contractile activity in the injured muscle due to biomechanical compensation of the contralateral limb. As a way of indirectly testing whether injured mice may have had altered muscle contractile activity during wheel running, maximal nerve evoked plantar flexor muscle strength was assessed (Fig. 5e-f). Contractile torque was substantially greater, although not completely restored, in injured muscle that underwent 4 weeks of voluntary wheel running compared to untreated VML-injured muscle indicating that 1) the injured muscle underwent functional loading during wheel running and 2) contractility of VML injured muscle is plastic. Together, these data suggest that the muscle remaining following VML injury is plastic in terms of contractility but not oxidative capacity.

**Figure 5:**
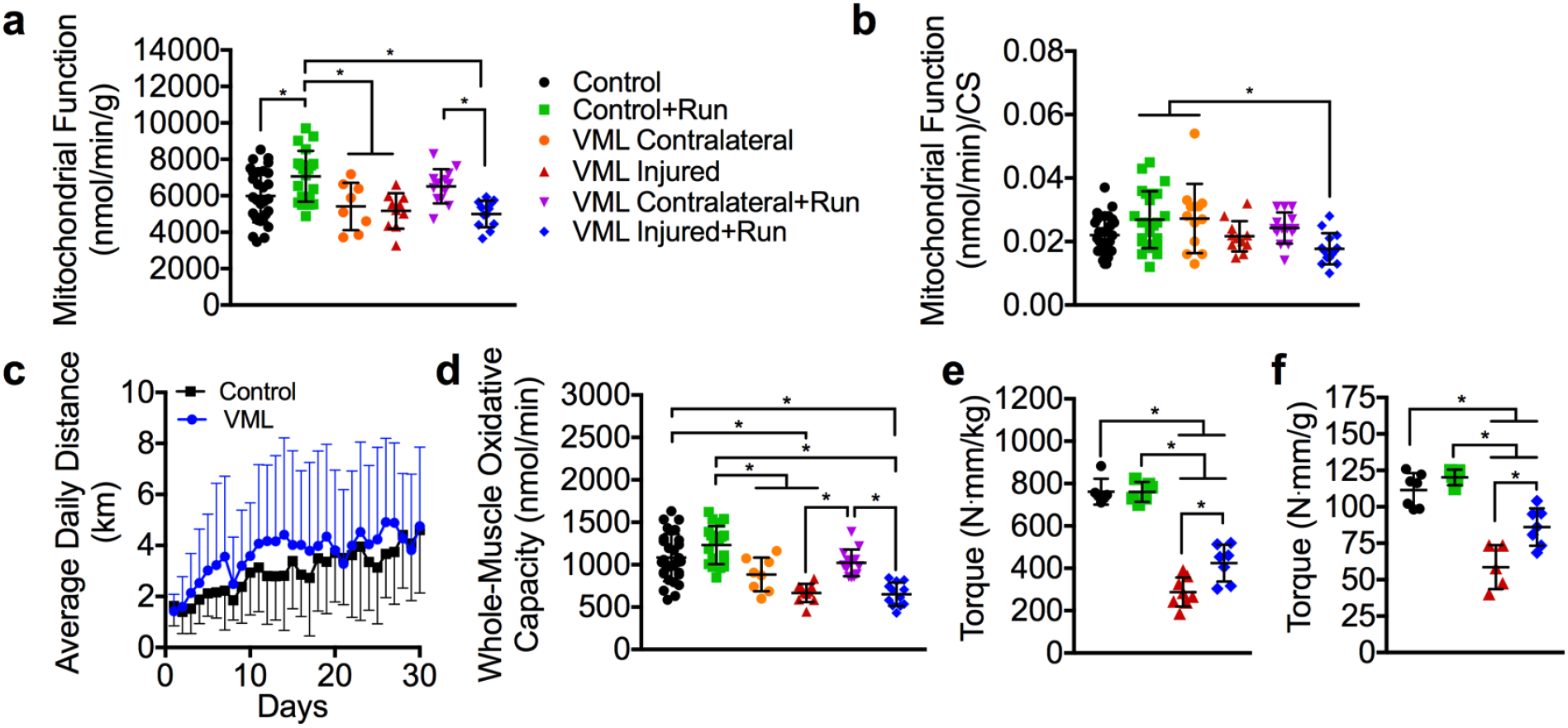
The effects of VML and voluntary wheel running on muscle strength and oxidative capacity 4 weeks post injury. (**a**) Mitochondrial respiratory function normalized by grams wet weight of permeabilized muscle fibers (n≥12 permeabilized fiber bundles from n=6 mice for each condition) did not adapt to exercise training in the VML injured limb. (**b**) Mitochondrial respiratory function normalized to citrate synthase enzyme activity (see Table 1). (**c**) Wheel running distance of control and VML injured mice. (**d**) Extrapolation of mitochondrial respiration rates (panel **a**) to entire muscle mass (see Table 1). Peak isometric torque normalized to (**e**) body mass and (**f**) plantarflexor mass in the VML injured limb was significantly greater after voluntary wheel running. Data analyzed by one-way ANOVA, *P<0.05. Error bars represent means ± SD.

### Vascular Disruptions are Likely not a Mechanism of Impaired Oxidative Plasticity

Next, we sought to elucidate a mechanism that could explain how VML injury results in impaired oxidative plasticity of the remaining muscle. We first considered that inadequate vascularization of the surrounding muscle tissue after injury might impede oxidative adaptations to exercise. VML likely results in severe disruptions to the vascular network adjacent to the injury area, which could disrupt delivery of oxygen and fuel to the mitochondria. Micro-CT imaging was employed to generate 3D models of the vascular network(45, 46) within injured and uninjured muscle of mice, both with and without voluntary wheel running therapy (Fig. 6a) to determine whether or not reductions in the vascular network in the muscle surrounding the injury contributes to the loss of oxidative plasticity. Interestingly, VML injured muscle, independent of wheel running therapy, had a significantly higher vascular volume compared to uninjured muscle indicating likely expansion of the vascular network after VML (Fig. 6b). While the viability of those vessels is unknown, it appears a reduction in the vascular network is not a limiting factor for oxidative capacity, prompting us to pursue another more viable hypothesis.

**Figure 6:**
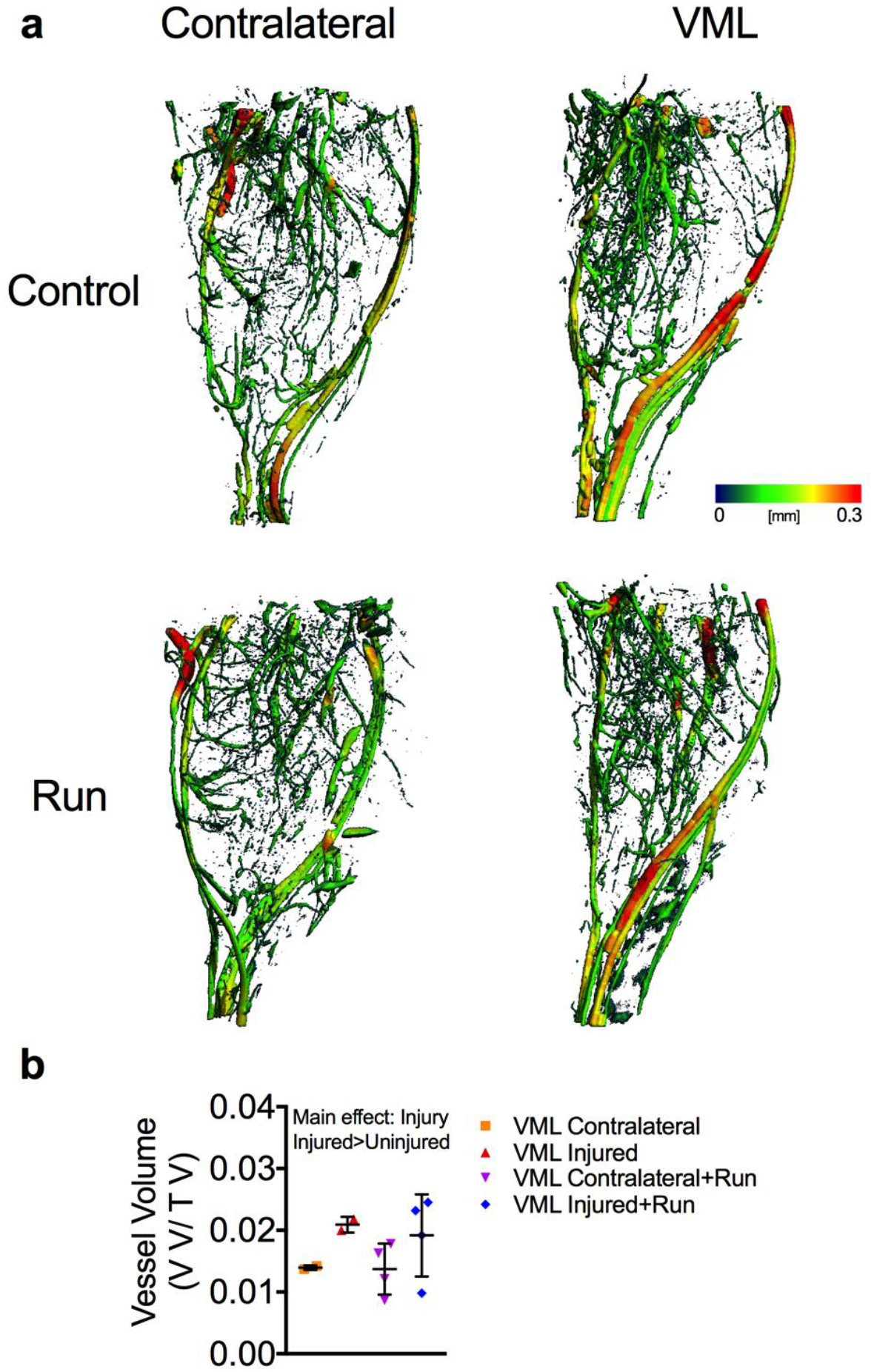
Changes in the vasculature networks is not altered following VML injury and wheel running. (**a**) Representative 3D reconstructions with vessel diameter mapping of vasculature in the posterior compartment of contralateral control and VML injured limbs. Additionally, mice were given access to running wheels (or remained sedentary) for one month following injury. (**b**) Quantification of blood vessel volume normalized to total volume. Data analyzed by two-way ANOVA; P=0.007 for main effect of muscle injury. Error bars represent means ± SD.

### Mitochondrial Biogenesis as a Mechanism for Impaired Oxidative Plasticity

Mitochondrial biogenesis is the cellular process of expanding the mitochondrial reticulum by the creation of new organelles and is responsive to cellular stimuli such as an increase in energy demand. Because mitochondrial biogenesis is a primary adaptation to exercise training and is necessary for exercise-induced increases in mitochondrial function and content, we thought that impaired mitochondrial biogenesis in VML injured muscle could potentially explain the lack of oxidative plasticity. To test this, we designed an acute stimulation protocol that would mimic a short bout of overload exercise capable of generating a stimulus for mitochondrial biogenesis(47–49). To control for potential differences in voluntary activation of the hindlimb between VML injured and control mice, direct *in vivo* stimulation of the sciatic nerve for complete activation the plantar flexor muscles was used. PGC-1α gene expression was measured after 30 minutes of unilateral stimulation on bi-laterally VML injured mice 14 days after injury as well as injury naïve mice. As expected, PGC-1α gene expression was ~3-fold greater in the stimulated limb of injury naïve mice compared to the unstimulated limb (Fig. 7a). However, in the stimulated limb of VML injured mice, PGC-1α gene expression did not increase after the stimulation indicating that mitochondrial biogenesis signaling was altered (Fig. 7a). This finding suggests that the lack of oxidative adaptations in VML injured tissue after therapeutic exercise is due to impairments in mitochondrial biogenesis signaling within the injured muscle.

**Figure 7:**
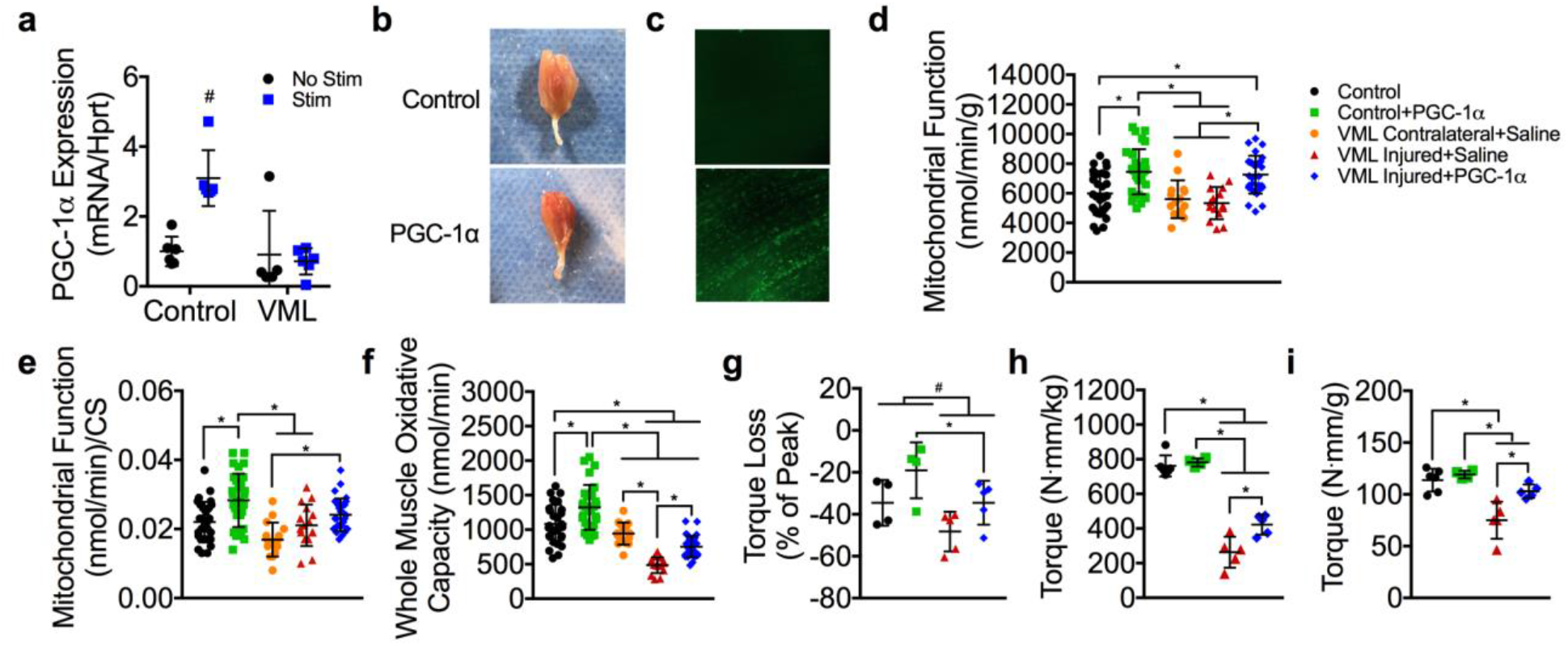
Effect of PGC-1α overexpression on oxidative capacity and plantarflexor muscle strength 4 weeks after VML injury. (**a**) PGC-1α gene expression 3 hours after completion of stimulation protocol (30 minutes of stimulation) for stimulated and non-stimulated limbs of control and VML injured mice; two-way ANOVA, interaction P<0.001, # indicates significantly different from all other experimental groups. (**b**) Representative images of whole gastrocnemius muscles with (bottom) and without (top) PGC-1α overexpression showing increased red hue in PGC-1α transfected muscle, indicating greater oxidation. (**c**) Representative images of muscle with and without GFP-PGC-1α transfection showing GFP fluorescence in PGC-1α transfected muscle. (**d**) Mitochondrial respiratory function normalized by grams wet weight of permeabilized muscle fibers (n≥15 permeabilized fiber bundles from n=5 mice for each condition). (**e**) Mitochondrial respiratory function normalized to citrate synthase enzyme activity. (**f**) Extrapolation of mitochondrial respiration rates (panel **d**) to entire muscle mass (see Table 1). (**g**) Plantarflexor torque loss following a fatiguing bout of 120 contractions; two-way ANOVA, P=0.014 for both main effects of PGC-1α (*) and group (#). Peak isometric torque normalized to (**h**) body mass and (**i**) plantarflexor mass was partially rescued in the VML injured limb after PGC-1α overexpression. Data analyzed by one-way ANOVA unless specified otherwise, *P<0.05. Throughout, error bars represent means ± SD.

### Overexpression of PGC-1α can Correct Oxidative Deficits and Partially Rescue Contractile Deficits in the Muscle Remaining after VML Injury

Our findings suggest that PGC-1α mediated mitochondrial biogenesis is a limiting factor that contributes to a lack of oxidative plasticity in the remaining muscle following VML injury. To explore this, we tested if bypassing contractile-dependent PGC-1α activation (i.e., electrical stimulation and wheel running) could rescue the oxidative capacity phenotype in the remaining muscle. Control and VML injured muscle were transfected with GFP-tagged PGC-1 plasmid driven by a CMV promoter by electroporation immediately after the VML injury(50) (Fig. 7b-c). Four weeks after transfection and injury, all mice, including VML-injured mice, had 25-35% greater mitochondrial function and content compared to non-transfected and vehicle (i.e. saline) transfected control mice indicated by state 3 respiration (function) and citrate synthase activity (content) (Fig. 7d-f, Table 1). Interestingly, the improvements in oxidative capacity were also concomitant with greater torque production and contractile fatigue resistance in VML-injured mice (Fig. 7g-h). In fact, PGC-1α overexpression rescued ~32% of the torque deficit in VML injured mice after accounting for plantar flexor muscle mass, although a ~10% deficit verses control still remained (Fig. 7i). These results highlight the potential benefit of developing treatment strategies targeted to restoration of mitochondrial biogenesis and by extension oxidative capacity in the muscle remaining after VML injury, as they may also be a useful therapeutic for restoring contractile function.

## Discussion

Current research on treatments for VML injury focus primarily on two areas: 1) designing physical bioengineered constructs with or without cellular and growth factor components to fill the void and facilitate endogenous or exogenous regeneration of lost muscle, and/or 2) designing structured physical therapy programs that strengthen the remaining muscle. Our study uncovered an important and novel aspect of VML pathophysiology, namely mitochondrial dysfunction, which is expected to provide critical contributions to both the development and evaluation of various treatment approaches for VML injury. Herein, we first identified mitochondrial network disorganization and dysfunction as a muscular complication (i.e., comorbidity) caused by VML injury. These complications, in addition to other comorbidities to injury, are expected to contribute to a hostile local and systemic environment that should be accounted for during the development and implementation of regenerative medicine approaches for VML. Second, we demonstrated that VML-injured muscle lacks the ability to oxidatively adapt to exercise, which is potentially a mechanism responsible for the limited efficacy of functional rehabilitation following VML.

One important characteristic of VML injury is the large loss in muscle contractile function for a relatively small loss in muscle tissue removed during the injury (see for reviw(9)). Similarly, in this study, there was a significant reduction in oxidative capacity acutely after VML, but it was unclear whether the loss in whole-muscle oxidative capacity was proportionate to the reduction in muscle volume. To explore this relationship, we generated an estimate of whole-muscle oxidative capacity, by extrapolating the oxygen consumption rate per mg of muscle to the entire muscle (Fig. 1e). This analysis revealed that the VML injured muscle had significantly greater reduction in oxidative capacity (~60%) compared to the amount of muscle tissue removed with VML (~10-15%; Fig. 1g). To determine if this relationship between oxidative capacity and muscle tissue volume occurs in other conditions of muscle loss, we performed an analysis of 15 studies spanning multiple conditions that are associated with a loss in muscle volume [i.e., denervation(27–31), aging(32–35), cachexia(36–38), immobilization(39), heart failure(40), ischemia reperfusion injury(41), and critical illness(42)]. On average, there was a ~29% loss in oxidative capacity after an average loss in muscle mass of ~36%, which suggests that oxidative capacity in VML injured muscle is disproportionately reduced in relation to the loss in muscle tissue. This finding is in line with the mitochondrial respiratory and structural deficits observed acutely after VML, and reveals the potential impact of the extensive mitochondrial damage throughout the remaining muscle.

The disproportionate reduction in oxidative capacity that is observed after VML is partially due to the mechanical damage from the injury itself, but mechanical damage alone cannot explain the extensive mitochondrial structural and functional abnormalities that occur after VML. Inflammation has been associated with mitochondrial dysfunction(51, 52) and an excessive inflammatory response, like the one that occurs after VML, could be a potential contributor to the mitochondrial damage in the remaining muscle. Generally, following a muscle injury, the inflammatory response is important for initiating endogenous recovery of muscle function and typically resolves within 3-7 days(53) depending on injury severity. However, following VML injury, the inflammation could potentially exacerbate the injury as it is heightened and prolonged, likely lasting for well more than a month(25, 54). The quantitative 2-photon scanning microscopy provided evidence that suggests the damaging effects of VML injury on mitochondrial organization and function extend well beyond the borders of the injury (Fig. 4). This widespread effect of VML injury could be due to the excessive inflammatory response that occurs after the injury, which would support the notion of an inflammation-based bystander injury in the myofibers surrounding the initially injured area(55, 56).

The mitochondria defects in VML-injured muscle may be an important contributor to the disproportionate loss in muscle function after VML. We expect that the extensive mitochondrial structural and functional disparities observed after VML creates an inhospitable environment to endogenous skeletal muscle regeneration, which perpetuates the chronic disproportionate loss in muscle function. This notion is supported by several lines of evidence that suggest mitochondria have an important role in skeletal muscle regeneration. First, mitochondrial biogenesis coincides with the time course of muscle regeneration(57, 58). Second, mitochondrial quality is necessary for successful regeneration, as Jash et al. demonstrated that restoring mitochondria with polycistronic RNAs after muscle injury can lead to greater activation and proliferation of endogenous satellite cell populations(59). Lastly, enhancing mitochondrial capacity is sufficient to accelerate recovery of muscle function suggesting that the functional quality of the mitochondrial network is important for the regeneration potential of the muscle(60). Given this evidence, it is not surprising that extensive damage to the mitochondrial network after VML would be associated with extreme loss in muscle function. Indeed, the data presented herein show that both muscle oxidative capacity and structure are reduced by similar magnitudes which supports the argument that damaged mitochondria in VML-injured muscle may be a major contributor to muscle dysfunction after VML.

The VML-injured muscle is an obvious candidate for rehabilitative therapy due to expected benefits of greater strength, fatigue resistance, and a healthier more favorable cellular environment for regenerative medicine approaches (e.g., biomaterial, stem cell, and growth factor-based therapies). However, VML-injured muscle does not only have compromised oxidative capacity, but also compromised oxidative plasticity, and physical rehabilitation interventions are pointless endeavors if the tissue is unable to remodel and adapt. Certainly, the parameters (i.e., optimal frequency, duration, and intensity) of post-VML rehabilitative care need to be validated, but identification of the mechanisms of limited plasticity in VML-injured muscle must be uncovered before specific and effective treatment regimens can be created.

PGC-1α gene expression did not increase with an acute bout of stimulation, which provides a potential mechanism for the impaired oxidative plasticity observed in VML-injured muscle. This phenomenon, the lack of response in PGC-1α gene expression, is likely the result of inadequate muscle activation during exercise. Damage to descending axons, intramuscular nerves, and neuromuscular junctions (NMJs) downstream of the directly activated sciatic nerve could impede complete muscle activation. Indeed, Beltran et al. reported that ~20% of VML patients suffer peripheral nerve injuries concurrently with the injury(61), and a recent report highlighted significant chronic motor neuron axotomy that occurs in VML-injured muscle(24). Furthermore, overexpression of PGC-1α induced greater oxidative capacity in the VML-injured muscle, which suggests that the mitochondrial biogenesis signaling pathway is intact and oxidative adaptations are possible if mitochondrial biogenesis is stimulated independent of contractile activity. These data support a framework in which VML-induced nerve damage likely limits muscle fiber activation and motor unit recruitment during exercise in the muscle tissue adjacent to the injury site, thus preventing oxidative adaptations.

Four weeks of PGC-1α overexpression in VML-injured muscle nearly completely rescued muscle strength normalized to muscle mass. These data indicate that PGC-1α plays an important role in recovery of muscle function after VML, but it remains to be seen exactly how PGC-1α overexpression mechanistically leads to greater strength. There are several possible ways that PGC-1α overexpression could improve contractility in VML-injured muscle. First, mitochondria have been shown to aid in muscle fiber sarcolemmal repair after injury(62, 63), meaning that myofibers with more mitochondria (i.e., PGC-1α overexpression) may be more resistant to cell death after injury. Second, PGC-1α overexpression has been shown to enhance structure and innervation of the NMJ, in particular the pre-synaptic integrity of the NMJ(4), which could partially compensate for neural damage caused by VML by facilitating retrograde neurotrophic signaling to motor neurons. Third, PGC-1α overexpression has been shown to prevent muscle atrophy, which is a common side effect of VML, by inhibiting FoxO3 signaling(64). Overall, our data show that PGC-1α overexpression and the resulting increases in oxidative capacity are associated with improved muscle strength after VML injury, but the exact mechanism or mechanisms by which PGC-1α overexpression causes greater muscle strength remains unclear and should be a focus of future research.

The improvements in functional capacity following PGC-1α overexpression are promising in that VML injured muscle was capable of oxidative adaptations and suggests that mitochondria may play an important and unaddressed role in the recovery of muscle strength after VML. Thus, future research and treatment strategies should be expanded to address limitations in oxidative adaptations to rehabilitation to potentially exploit the role of mitochondria in recovery of muscle function after VML. In conclusion, this work uncovers a novel element of VML pathophysiology and provides valuable insight for development and optimization of future VML treatment strategies.

## Methods

### Experimental Design

Male C57BL/6 mice were housed at 20-23°C on a 12:12-hr light-dark cycle, with food and water provided ad libitum. At the time of randomization to experimental groups all mice were 9 weeks of age. For the acute study, (n=8) mice were randomized to unilateral VML injury for either 3 or 7 days. For the wheel running study, (n=5-7 per group) mice were randomized to uninjured control and 4 weeks of wheel running (Control+Run), unilateral VML alone (VML Contralateral; VML Injured), or unilateral VML and 4 weeks of wheel running (VML Contralateral+Run; VML Injured+Run). Mice within running groups were given access to voluntary running wheels 72 hours after VML injury. For the PGC-1α transfection study, (n=4-5 per group) mice were randomized to uninjured control and PGC-1α plasmid (Control+PGC-1α), unilateral VML and empty vector (i.e. saline) (VML Contralateral+Saline; VML Injured+Saline), or bilateral VML and PGC-1α plasmid (VML Injured+PGC-1α). All outcome measures were also collected in an uninjured, untreated control group (Control, n=7) and was used as a reference group across studies. Plantarflexor peak isometric strength and gastrocnemius mitochondrial respiration were measured after 3 and 7 days (acute study), after 4 weeks of voluntary running (wheel running study), and after 4 weeks of PGC-1α overexpression (PGC-1α transfection study). Plantarflexor fatigability was also measured after 4 weeks of PGC-1α overexpression (PGC-1α transfection study). Immediately after assessment of peak strength, the gastrocnemius muscle was harvested and prepared for mitochondrial respiration measurements and mitochondrial enzyme assays. All procedures were approved and performed in accordance with relevant guidelines and regulations by the Institutional Animal Care and Use Committee at the University of Georiga.

### Surgical Creation of VML Injury

VML injury was conducted on the posterior compartment of anesthetized (isoflurane 1.5–2.0%) mice. All mice received administration of buprenorphine-SR (1.2 mg/kg; s.c.) for pain management 30 minutes prior to surgery. A posterior incision was made to expose the posterior compartment muscles, and blunt dissection was used to remove the fascia and hamstrings to expose the gastrocnemius muscle. A small metal plate was inserted behind the gastrocnemius and soleus muscles and a 4-mm biopsy punch was used to remove 22.1±2.9 mg of muscle volume (~10-15% of uninjured gastrocnemius mass) from the center of the gastrocnemius muscle. In a subset of mice a TA muscle VML injury was made for evaluation of 2-photon imaging of mitochondrial structure. Under the same induction and pain management techniques, an anterior incision was made to expose the TA muscle and the fascia was removed. A 3-mm biopsy punch was used to surgically create the VML injury in the middle of the muscle (7.46±1.2 mg removed). For both procedures, following the VML, the skin incision was sutured closed (6-0 silk). There were no adverse events noted in any of the experimental groups.

### Voluntary Wheel Running

Injured and control mice in the one month wheel running study, were housed individually and given free access to a running wheel (Columbus Instruments, Columbus, Ohio). Sedentary VML mice were housed in a standard mouse cage without access to a running wheel. Daily running totals were calculated from wheel revolutions collected at 5 min intervals and are presented as a daily average of distance ran.

### *In Vivo* Muscle Function

Prior to assessment of *in vivo* peak isometric torque of the ankle plantarflexors, mice were anaesthetized using 1.5% isoflurane at an oxygen flow rate of 0.4L/min. The left hindlimb was depilated and aseptically prepared and the foot placed in a foot-plate attached to a servomotor (Model 300C-LR; Aurora Scientific, Aurora, Ontario, Canada). The left peroneal nerve was severed and platinum-iridium needle electrodes (Model E2-12; Grass Technologies, West Warwick, RI) were placed on either side of the sciatic nerve to elicit isolated contraction of the plantarflexor muscles. Peak isometric torque was defined as the greatest torque measured during a 200-ms stimulation using 1-ms square-wave pulses at 300 Hz and increasing amperage 0.6 to 2.0 mA (models S48 and SIU5; Grass Technologies). Fatigability of the plantarflexors muscles was assessed using 120 submaximal isometric contractions were performed in 2 min using 330 ms stimulations at 50 Hz.

### Stimulation of Exercise Bout

To assess the integrity of mitochondrial biogenesis signaling in VML injured muscle, a 30 minute *in vivo* electrical stimulation protocol was used to simulate an acute bout of exercise. The stimulation protocol was performed two weeks after VML injury on the left limb of age-matched control mice (n=6) and bilateral VML injured mice (n=6). Electrical stimulation was used instead of voluntary exercise to ensure activation of the injured gastrocnemius muscles. Prior to the start of the stimulation protocol, platinum-iridium needle electrodes were placed around the sciatic nerve of anesthetized (isoflurane 1.5-2.0%) mice and electrical current was optimized for peak torque generation. The electrical stimulation protocol consisted of 10 sets of 1800 contractions (parameters: pulse frequency=100, pulse width=0.1, pulses per train=1, train frequency=10 Hz) conducted over 30 minutes. At the end of the protocol, gastrocnemius muscles were quickly harvested, flash frozen in liquid nitrogen, and stored at −80°C for later qRT-PCR analysis.

### Gene Expression

RNA was isolated from frozen gastrocnemius muscles using an RNeasy kit (QIAGEN) and cDNA was generated using a High Capacity cDNA Reverse Transcription Kit (Applied Biosystems). iQ SYBR Green Supermix (Bio-Rad) and the following sequence-specific primer was used to assess mRNA levels for *PGC-1α*, (For: 5’-AGC CGT GAC CAC TGA CAA CGA G-3’; Rev: 5’-GCT GCA TGG TTC TGA GTG CTA AG-3’). NormFinder Software(65) was used to identify the most stable reference gene between 18s, Hprt, and Hsp90. Hprt (For: 5’-TCAACGGGGGACATAAAAGT-3’; Rev: 5’-TGCATTGTTTTACCAGTGTCAA-3’) was identified as the most stable gene in this VML model and therefore was the reference gene of choice for this analysis. Relative gene expression was calculated using the 2^−ΔΔCT^ method.

### Mitochondrial Assays

Immediately following sacrifice, the medial and lateral gastrocnemius muscles from uninjured and injured limbs were dissected on a chilled aluminum block in 4°C buffer X containing 7.23mM K_2_EGTA, 2.77mM Ca K_2_EGTA, 20mM imidazole, 20mM taurine, 5.7mM ATP, 14.3mM PCr, 6.56mM MgCl_2_-6H_2_O, 50mM k-MES. Muscles were carefully dissected <1mg bundles of muscle fibers as reported by Kuznetsov et al.(66). Fiber bundles were permeabilized via an incubation (i.e. rocking) in buffer X and saponin (50 μg/ml) at 4°C for 30 minutes. Following permeabilization, muscle fiber bundles were rinsed for 15 minutes in buffer Z (105mM k-MES, 30mM KCl, 10mM KH_2_PO_4_, 5mM MgCl_2_, 0.5 mg/ml BSA, 1mM EGTA) at 4°C. All respiration measurements were performed using a Clark-type electrode (Oxygraph Plus System, Hansatech Instruments, UK) at 25°C. Prior to each experiment, the electrode was calibrated according to the manufacturer’s instructions and 1 ml of oxygen infused buffer Z was added to the chamber. Muscle fiber bundles were weighed (~2.5 mg for all samples) and added to the chamber. State 4 respiration (leak respiration in the absence of ADP) was initiated by the addition of glutamate (10mM) and malate (5mM). State 3 respiration (respiration coupled to ATP synthesis) was initiated by the addition of ADP (2.5mM) and succinate (10mM). Cytochrome *c* (10μM) was added to measure the integrity of the outer mitochondrial membrane (data not shown). Respiration rates were expressed relative to the mg of tissue loaded into each oxygraph chamber as well as to citrate synthase activity to account for differences in mitochondrial content between samples.

Citrate synthase activity was measured using a protocol modified from Srere(67), Briefly, ~20 mg of gastrocnemius muscle was homogenized in ~800 μl of 33mM phosphate buffer (pH 7.0). 5 μl of homogenate, 173.74 μl of 100mM Tris buffer (pH 8.0), 17.51 μl of DTNB, 8.75 μl of acetyl CoA, and 20 μl of oxaloacetate were combined in a well of a 96 well plate. Absorbance was measured at 405nm every 10 seconds for 3 minutes and citrate synthase activity was determined from the change in optical density over that time. Enzyme activities were normalized to mg of tissue in the sample homogenate.

### Micro-CT Angiography

A subset of C57Bl/6 mice were randomly assigned to either VML (VML Contralateral; VML Injured) or VML and 4 weeks of wheel running (VML Contralateral+Run; VML Injured+Run). Unilateral VML injury was performed on the posterior compartment of all mice. Four weeks after injury, micro-CT angiography was used to quantitatively evaluate hindlimb vasculature(45, 46). After animal euthanasia, the vasculature was cleared with 0.9% saline, perfusion fixed with 10% neutral buffered formalin, rinsed again with saline, and injected with Microfil contrast agent (MV-122, Flow Tech Inc.). Samples were stored at 4°C overnight to allow for polymerization of the contrast agent. Hind limbs were harvested and stored in PBS at 4°C until imaging.

For imaging, samples were oriented with long axis of the tibia extending in the z-direction for micro-CT scanning (μCT50, Scanco Medical). Scans were performed on the lower leg with an applied electric potential of 55 kVp, a current of 145 μA, and an isometric voxel size of 20 μm. After automated reconstruction to 2D slice tomograms, contouring was performed on slices to mark a total muscle volume of analysis that excluded bones and only selected the musculature in the posterior compartment of the lower hindlimb. A global X-ray attenuation threshold was applied for segmentation of Microfil perfused vasculature, and a Gaussian low-pass filter was used for smoothing and noise suppression. This produced 3D images and volumetric quantifications (using direct distance transformation methods included in Scanco software) for vascular anatomy with the outcome measure of vascular volume normalized to total volume. All investigators involved in scanning and analysis were blinded to experimental groups.

### Plasmid and Transfection

Electroporation and PGC-1α plasmid transfection were conducted at the time of VML injury. GFP-PGC1 plasmid expressing eGFP-tagged mouse PGC1a was acquired from Addgene(50). For *in vivo* electroporation, GFP-PGC1 plasmid was prepared by cesium chloride density-gradient centrifugation and isopropanol precipitation as previous reported(68). *In vivo* electroporation of mouse gastrocnemius muscles was performed as described by Aihara et al.(69). Briefly, 20 μl of GFP-PGC1 plasmid (concentration=2.8 μg/μL) was injected at 2 sites: medial and lateral gastrocnemius muscles. Electroporation was conducted with a BTX ECM 830 electroporation system equipped with 5 mm 2-needle arrays. The following settings of the electronic pulses were use: LV= 00V/99 msec, set voltage=100 V, set pulse length=50 msec, set number of pulses=3 pulses. When 3 pulses were done, the 2-needle array was reversed, and 3 addition pulses were applied to the muscle with the above settings.

### 2-Photon Scanning Microscopy

Unilateral VML injury was performed on the left TA muscle of C57Bl/6 mice ubiquitously expressing mitochondrial Dendra2 green/red photoswitchable monomeric fluorescent protein (Jackson Laboratory, #018385). We elected to use the TA for the imaging rather than the gastrocnemius muscle used for other studies herein, because of its accessibility and lack of pennation differences across medial and lateral aspects of the muscle. Imaging was performed immediately (data not shown), 3, 7, and 28 days after VML injury, and the contralateral limb was imaged at each timepoint as a control.

Prior to imaging, the TA was extracted, placed in buffer X (*see Methods: Mitochondrial Assays*), and secured to a dissection gel with pins. 2-photon microscopy was used to characterize the mitochondrial network of the muscle fibers remaining after VML. We used a Ti:Sapphire laser (Coherent Chameleon Ultra II), with 840nm and 940nm wavelength and 130 fs pulses duration for excitation of the Dendra2 fluorescent protein, with an NA = 1.1 objective lens (Olympus LUMFLN 60XW) and a 509/22nm filter for fluorescence collection. For the 3, 7, and 28-day time points, we imaged within an area of approximately 1.0 mm^2^ in the proximity of the injury site. Within each area, we collected multiple z-stacks (n>10), with resolution and pixel size small enough to satisfy the Nyquist sampling theorem (dx = dy = 0.3125 μm, dz = 1 μm). Additional z-stacks (n=3-6) were collected proximal to the border of the injury at increasing distances (0mm, 0.5mm, 1.5mm, 2.0mm, and 2.5mm) away from the injury site toward the origin of the muscle.

To analyze the mitochondrial network organization, we developed an angular Fourier filtering (AFF) method, based on a Fourier metric previously used for sensorless Adaptive Optics (70). Transforming an image from the spatial domain (fig. S1 A,C) to the Fourier domain (fig. S1 B,D), converts it to an array of weighted coefficients that include information such as periodicity and angle of the features in the image (Fig. 8). Our method uses a wedge filter ψ with a gaussian profile w.r.t. φ in cylindrical coordinates ρ,φ, (figure S1 E) to collect information about features at an angle α:

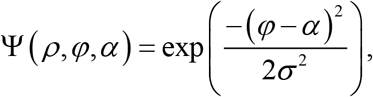

**Figure 8:**
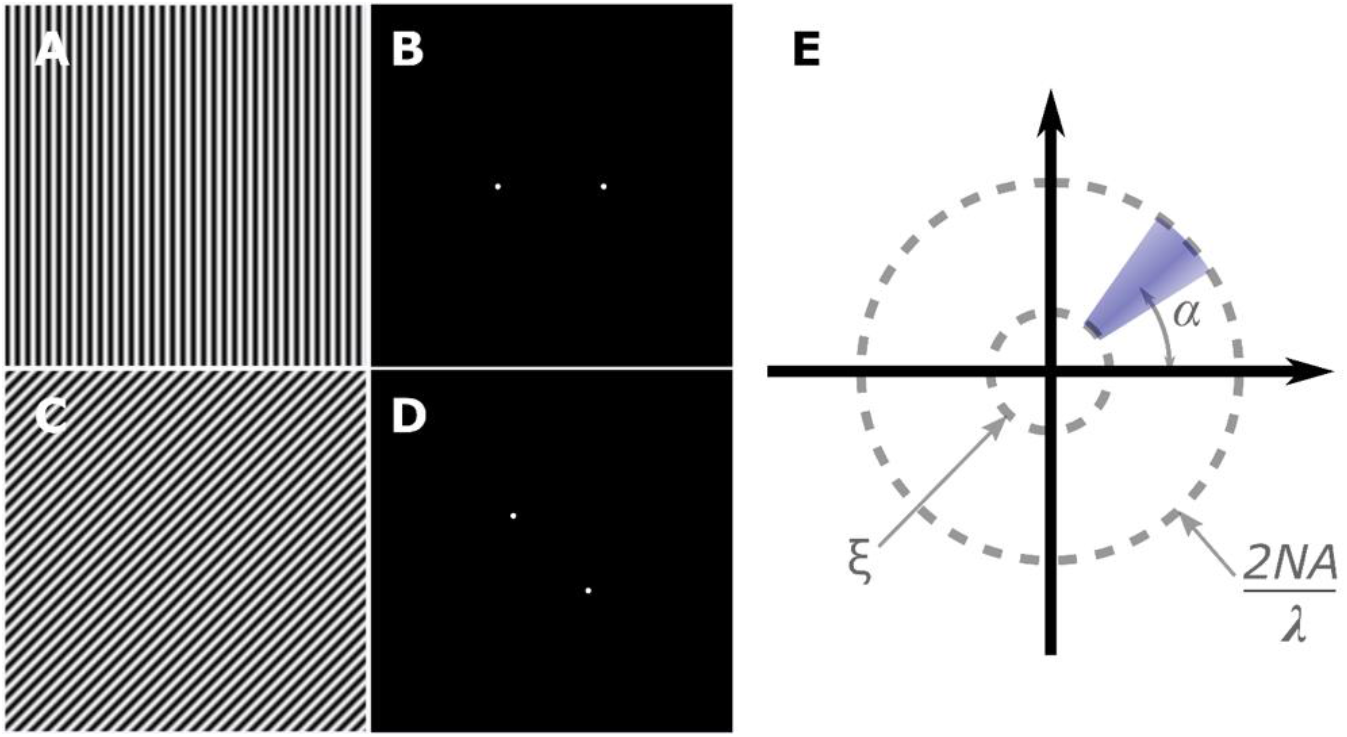
Illustration of the angular Fourier filter. A and C show sinusoidal patterns in the spatial domain with frequency of ω at 0° and 45°, respectively. Corresponding power spectral density (PSD) of A and C are shown in B and D. In the PSD there exists only two peaks at the frequencies ω and −ω, with zero frequency being at the center. Because the angle of the pattern in the spatial domain image (A,C) are followed in the PSD (B,D) we can use this approach to characterize the angular distribution of patterned structures such as the mitochondrial network. Our Angular Fourier Filter is shown in E, depicting the gaussian wedge filter placed at angle α. It also acts as a bandpass filter to suppress features smaller than the diffraction limit to avoid unwanted noise, and larger than the mitochondrial network to avoid distortion of the analysis.

Where σ is the standard deviation of gaussian filter. We then calculate the 2D Fourier transform of each image *i* at depth *z*:

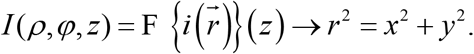

We rotate the filter from 0 to 180 degrees for each optically sectioned image at depth z to produce a histogram of the angle of alignment and periodicity of the structure. Each point in the generated 2D histogram AFF is calculated using:

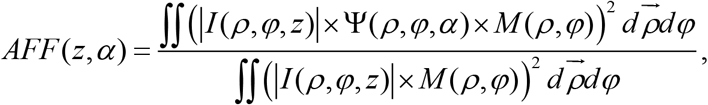

where M is a mask determined by the numerical aperture of the microscope *NA*,the wavelength of the emitted light *λ*, and the frequency lower bound *ξ* (to suppress features larger than twice the mitochondrial network period):

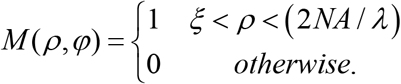

AFF is an ideal analysis for assessing mitochondrial structural organization because it can detect the 2 dominant angles of alignment in uninjured mitochondrial networks: 1) mitochondrial network alignment parallel to muscle fiber orientation and 2) mitochondrial network alignment perpendicular to muscle fiber orientation. AFF was used in each frame from a z-stack to produce 3D mesh plots (Fig. 2). The magnitude of the peaks within the 3D mesh plots represents the strength of alignment of mitochondrial structures to a particular angle. To quantify the organization of mitochondrial structures the ratio of peak alignment strength of a given z-stack and its average alignment strength across the entire z-stack was calculated (Fig. 3c & d), which we call the alignment ratio (AR):

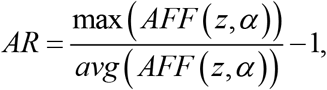

where max and avg are functions returning the maximum and the average of the arrays, respectively.

### Statistical Analysis

Data are presented in the results as mean ± SD and represented graphically as dot plots. A multi-factor repeated measures analysis of variance (ANOVA) was used to analyze assessments wheel running performance (injury group by time, repeated). A multi-factor repeated measures ANOVA was also used to analyze data from the acute VML study (time post injury by experimental limb). Gene expression data was analyzed using nonparametric tests in REST 2009 Software (M. Pfaffl, Technical University Munich, and QIAGEN). A two-way ANOVA was used to analyze torque as well as fatigue resistance after PGC-1α transfection (group by treatment). All other data were analyzed using one-way ANOVA. All data were required to pass normality (Shapiro-Wilk) and equal variance tests (Brown-Forsythe *F* test) before proceeding with the ANOVA. Differences among groups are only reported where significant interactions were observed and subsequently tested with Tukey’s *post hoc* test using JMP statistical software (version 13, SAS, Cary, NC). Group main effects are reported where significant interactions were not observed. An α level of 0.05 was used for all analyses.

## Data Availability

The datasets used and/or analyzed during the current study are primarily presented in the current manuscript and are available from the corresponding author on reasonable request. The executable code used to analyze mitochondrial network organization is available through github (link pending).

## Acknowledgements

This work was supported by funding from: the Alliance for Regenerative Rehabilitation Research & Training (AR3T) awarded to SMG and JAC, which is supported by the Eunice Kennedy Shriver National Institute of Child Health and Human Development (NICHD), National Institute of Neurological Disorders and Stroke (NINDS), and National Institute of Biomedical Imaging and Bioengineering (NIBIB) of the National Institutes of Health under Award Number P2CHD086843; The Assistant Secretary of Defense for Health Affairs endorsed by the Department of Defense, through the Clinical & Rehabilitative Medicine Research Program, FY17 Neuromusculoskeletal Injuries Rehabilitation Research Award (W81XWH-18-1-0710 to SMG and JAC). Opinions, interpretations, conclusions and recommendations are those of the authors and are not necessarily endorsed by the Department of Defense or National Institutes of Health

## Author Contributions

WMS, SMG, and JAC designed the study

WMS, ASN, KFT, MJM, LK, AEQ, NTJ, LJM, HY, AY, REG, SMG, and JAC performed experiments and collected data

WMS, ASN, KFT, MJM, LK, AEQ, NTJ, LJM, HY, AY, REG, SMG, and JAC analyzed the data

WMS, SMG, and JAC wrote the manuscript; all authors have read and approved the final version of this manuscript

## References

1. Gollnick PD, Armstrong RB, Saltin B, Saubert CWt, Sembrowich WL, Shepherd RE. Effect of training on enzyme activity and fiber composition of human skeletal muscle. J Appl Physiol. 1973;34(1):107–11. Epub 1973/01/01. doi: 10.1152/jappl.1973.34.1.107. PubMed PMID: 4348914.

2. Ingjer F. Effects of endurance training on muscle fibre ATP-ase activity, capillary supply and mitochondrial content in man. J Physiol. 1979;294:419–32. Epub 1979/09/01. PubMed PMID: 159945; PMCID: PMC1280565.

3. Hood DA, Tryon LD, Carter HN, Kim Y, Chen CC. Unravelling the mechanisms regulating muscle mitochondrial biogenesis. Biochem J. 2016;473(15):2295–314. Epub 2016/07/30. doi: 10.1042/BCJ20160009. PubMed PMID: 27470593.

4. Arnold AS, Gill J, Christe M, Ruiz R, McGuirk S, St-Pierre J, Tabares L, Handschin C. Morphological and functional remodelling of the neuromuscular junction by skeletal muscle PGC-1alpha. Nat Commun. 2014;5:3569. Epub 2014/04/02. doi: 10.1038/ncomms4569. PubMed PMID: 24686533; PMCID: PMC4846352.

5. Fernandez-Marcos PJ, Auwerx J. Regulation of PGC-1alpha, a nodal regulator of mitochondrial biogenesis. Am J Clin Nutr. 2011;93(4):884S–90. Epub 2011/02/04. doi: 10.3945/ajcn.110.001917. PubMed PMID: 21289221; PMCID: PMC3057551.

6. Rowe GC, Raghuram S, Jang C, Nagy JA, Patten IS, Goyal A, Chan MC, Liu LX, Jiang A, Spokes KC, Beeler D, Dvorak H, Aird WC, Arany Z. PGC-1alpha induces SPP1 to activate macrophages and orchestrate functional angiogenesis in skeletal muscle. Circ Res. 2014;115(5):504–17. Epub 2014/07/11. doi: 10.1161/CIRCRESAHA.115.303829. PubMed PMID: 25009290; PMCID: PMC4524357.

7. Egan B, Zierath JR. Exercise metabolism and the molecular regulation of skeletal muscle adaptation. Cell Metab. 2013;17(2):162–84. Epub 2013/02/12. doi: 10.1016/j.cmet.2012.12.012. PubMed PMID: 23395166.

8. Haskell WL, Lee IM, Pate RR, Powell KE, Blair SN, Franklin BA, Macera CA, Heath GW, Thompson PD, Bauman A. Physical activity and public health: updated recommendation for adults from the American College of Sports Medicine and the American Heart Association. Med Sci Sports Exerc. 2007;39(8):1423–34. Epub 2007/09/01. doi: 10.1249/mss.0b013e3180616b27. PubMed PMID: 17762377.

9. Corona BT, Wenke JC, Ward CL. Pathophysiology of Volumetric Muscle Loss Injury. Cells Tissues Organs. 2016;202(3-4):180–8. Epub 2016/11/09. doi: 10.1159/000443925. PubMed PMID: 27825160.

10. Corona BT, Rivera JC, Owens JG, Wenke JC, Rathbone CR. Volumetric muscle loss leads to permanent disability following extremity trauma. J Rehabil Res Dev. 2015;52(7):785–92. Epub 2016/01/09. doi: 10.1682/JRRD.2014.07.0165. PubMed PMID: 26745661.

11. Aurora A, Garg K, Corona BT, Walters TJ. Physical rehabilitation improves muscle function following volumetric muscle loss injury. BMC Sports Sci Med Rehabil. 2014;6(1):41. Epub 2015/01/20. doi: 10.1186/2052-1847-6-41. PubMed PMID: 25598983; PMCID: PMC4297368.

12. Aurora A, Roe JL, Corona BT, Walters TJ. An acellular biologic scaffold does not regenerate appreciable de novo muscle tissue in rat models of volumetric muscle loss injury. Biomaterials. 2015;67:393–407. Epub 2015/08/11. doi: 10.1016/j.biomaterials.2015.07.040. PubMed PMID: 26256250.

13. Corona BT, Garg K, Ward CL, McDaniel JS, Walters TJ, Rathbone CR. Autologous minced muscle grafts: a tissue engineering therapy for the volumetric loss of skeletal muscle. Am J Physiol Cell Physiol. 2013;305(7):C761–75. Epub 2013/07/26. doi: 10.1152/ajpcell.00189.2013. PubMed PMID: 23885064.

14. Quarta M, Cromie M, Chacon R, Blonigan J, Garcia V, Akimenko I, Hamer M, Paine P, Stok M, Shrager JB, Rando TA. Bioengineered constructs combined with exercise enhance stem cell-mediated treatment of volumetric muscle loss. Nat Commun. 2017;8:15613. Epub 2017/06/21. doi: 10.1038/ncomms15613. PubMed PMID: 28631758; PMCID: PMC5481841.

15. Greising SM, Warren GL, Southern WM, Nichenko AS, Qualls AE, Corona BT, Call JA. Early rehabilitation for volumetric muscle loss injury augments endogenous regenerative aspects of muscle strength and oxidative capacity. BMC Musculoskelet Disord. 2018;19(1):173. Epub 2018/05/31. doi: 10.1186/s12891-018-2095-6. PubMed PMID: 29843673; PMCID: PMC5975473.

16. Garg K, Ward CL, Hurtgen BJ, Wilken JM, Stinner DJ, Wenke JC, Owens JG, Corona BT. Volumetric muscle loss: persistent functional deficits beyond frank loss of tissue. J Orthop Res. 2015;33(1):40–6. Epub 2014/09/19. doi: 10.1002/jor.22730. PubMed PMID: 25231205.

17. Dziki J, Badylak S, Yabroudi M, Sicari B, Ambrosio F, Stearns K, Turner N, Wyse A, Boninger ML, Brown EHP, Rubin JP. An acellular biologic scaffold treatment for volumetric muscle loss: results of a 13-patient cohort study. NPJ Regen Med. 2016;1:16008. Epub 2016/07/21. doi: 10.1038/npjregenmed.2016.8. PubMed PMID: 29302336; PMCID: PMC5744714.

18. Mase VJ, Jr., Hsu JR, Wolf SE, Wenke JC, Baer DG, Owens J, Badylak SF, Walters TJ. Clinical application of an acellular biologic scaffold for surgical repair of a large, traumatic quadriceps femoris muscle defect. Orthopedics. 2010;33(7):511. Epub 2010/07/09. doi: 10.3928/01477447-20100526-24. PubMed PMID: 20608620.

19. Lowell BB, Shulman GI. Mitochondrial dysfunction and type 2 diabetes. Science. 2005;307(5708):384–7. Epub 2005/01/22. doi: 10.1126/science.1104343. PubMed PMID: 15662004.

20. Madamanchi NR, Runge MS. Mitochondrial dysfunction in atherosclerosis. Circ Res. 2007;100(4):460–73. Epub 2007/03/03. doi: 10.1161/01.RES.0000258450.44413.96. PubMed PMID: 17332437.

21. Glancy B, Hartnell LM, Combs CA, Femnou A, Sun J, Murphy E, Subramaniam S, Balaban RS. Power Grid Protection of the Muscle Mitochondrial Reticulum. Cell Rep. 2017;19(3):487–96. Epub 2017/04/20. doi: 10.1016/j.celrep.2017.03.063. PubMed PMID: 28423313; PMCID: PMC5490369.

22. Glancy B, Hartnell LM, Malide D, Yu ZX, Combs CA, Connelly PS, Subramaniam S, Balaban RS. Mitochondrial reticulum for cellular energy distribution in muscle. Nature. 2015;523(7562):617–20. Epub 2015/08/01. doi: 10.1038/nature14614. PubMed PMID: 26223627.

23. Greising SM, Dearth CL, Corona BT. Regenerative and Rehabilitative Medicine: A Necessary Synergy for Functional Recovery from Volumetric Muscle Loss Injury. Cells Tissues Organs. 2016;202(3-4):237–49. Epub 2016/11/09. doi: 10.1159/000444673. PubMed PMID: 27825146; PMCID: PMC5553044.

24. Corona BT, Flanagan KE, Brininger CM, Goldman SM, Call JA, Greising SM. Impact of volumetric muscle loss injury on persistent motoneuron axotomy. Muscle Nerve. 2018;57(5):799–807. Epub 2017/11/17. doi: 10.1002/mus.26016. PubMed PMID: 29144551.

25. Aguilar CA, Greising SM, Watts A, Goldman SM, Peragallo C, Zook C, Larouche J, Corona BT. Multiscale analysis of a regenerative therapy for treatment of volumetric muscle loss injury. Cell Death Discov. 2018;4:33. Epub 2018/03/14. doi: 10.1038/s41420-018-0027-8. PubMed PMID: 29531830; PMCID: PMC5841404.

26. Goldman SM, Henderson BEP, Walters TJ, Corona BT. Co-delivery of a laminin-111 supplemented hyaluronic acid based hydrogel with minced muscle graft in the treatment of volumetric muscle loss injury. PLoS ONE. 2018;13(1):e0191245. Epub 2018/01/13. doi: 10.1371/journal.pone.0191245. PubMed PMID: 29329332; PMCID: PMC5766229.

27. Adhihetty PJ, O'Leary MF, Chabi B, Wicks KL, Hood DA. Effect of denervation on mitochondrially mediated apoptosis in skeletal muscle. J Appl Physiol (1985). 2007;102(3):1143–51. Epub 2006/11/24. doi: 10.1152/japplphysiol.00768.2006. PubMed PMID: 17122379.

28. Kuo YT, Shih PH, Kao SH, Yeh GC, Lee HM. Pyrroloquinoline Quinone Resists Denervation-Induced Skeletal Muscle Atrophy by Activating PGC-1alpha and Integrating Mitochondrial Electron Transport Chain Complexes. PLoS ONE. 2015;10(12):e0143600. Epub 2015/12/10. doi: 10.1371/journal.pone.0143600. PubMed PMID: 26646764; PMCID: PMC4672922.

29. O'Leary MF, Vainshtein A, Carter HN, Zhang Y, Hood DA. Denervation-induced mitochondrial dysfunction and autophagy in skeletal muscle of apoptosis-deficient animals. Am J Physiol Cell Physiol. 2012;303(4):C447–54. Epub 2012/06/08. doi: 10.1152/ajpcell.00451.2011. PubMed PMID: 22673615.

30. Singh K, Hood DA. Effect of denervation-induced muscle disuse on mitochondrial protein import. Am J Physiol Cell Physiol. 2011;300(1):C138–45. Epub 2010/10/15. doi: 10.1152/ajpcell.00181.2010. PubMed PMID: 20943961.

31. Tryon LD, Crilly MJ, Hood DA. Effect of denervation on the regulation of mitochondrial transcription factor A expression in skeletal muscle. Am J Physiol Cell Physiol. 2015;309(4):C228–38. Epub 2015/06/13. doi: 10.1152/ajpcell.00266.2014. PubMed PMID: 26063705; PMCID: PMC4537931.

32. Joseph AM, Adhihetty PJ, Buford TW, Wohlgemuth SE, Lees HA, Nguyen LM, Aranda JM, Sandesara BD, Pahor M, Manini TM, Marzetti E, Leeuwenburgh C. The impact of aging on mitochondrial function and biogenesis pathways in skeletal muscle of sedentary high- and low-functioning elderly individuals. Aging Cell. 2012;11(5):801–9. Epub 2012/06/12. doi: 10.1111/j.1474-9726.2012.00844.x. PubMed PMID: 22681576; PMCID: PMC3444680.

33. Kang C, Chung E, Diffee G, Ji LL. Exercise training attenuates aging-associated mitochondrial dysfunction in rat skeletal muscle: role of PGC-1alpha. Exp Gerontol. 2013;48(11):1343–50. Epub 2013/09/03. doi: 10.1016/j.exger.2013.08.004. PubMed PMID: 23994518.

34. Picard M, Ritchie D, Thomas MM, Wright KJ, Hepple RT. Alterations in intrinsic mitochondrial function with aging are fiber type-specific and do not explain differential atrophy between muscles. Aging Cell. 2011;10(6):1047–55. Epub 2011/09/22. doi: 10.1111/j.1474-9726.2011.00745.x. PubMed PMID: 21933339.

35. Barnouin Y, McPhee JS, Butler-Browne G, Bosutti A, De Vito G, Jones DA, Narici M, Behin A, Hogrel JY, Degens H. Coupling between skeletal muscle fiber size and capillarization is maintained during healthy aging. J Cachexia Sarcopenia Muscle. 2017;8(4):647–59. Epub 2017/04/07. doi: 10.1002/jcsm.12194. PubMed PMID: 28382740; PMCID: PMC5566646.

36. Brown JL, Rosa-Caldwell ME, Lee DE, Blackwell TA, Brown LA, Perry RA, Haynie WS, Hardee JP, Carson JA, Wiggs MP, Washington TA, Greene NP. Mitochondrial degeneration precedes the development of muscle atrophy in progression of cancer cachexia in tumour-bearing mice. J Cachexia Sarcopenia Muscle. 2017;8(6):926–38. Epub 2017/08/29. doi: 10.1002/jcsm.12232. PubMed PMID: 28845591; PMCID: PMC5700433.

37. Fermoselle C, Garcia-Arumi E, Puig-Vilanova E, Andreu AL, Urtreger AJ, de Kier Joffe ED, Tejedor A, Puente-Maestu L, Barreiro E. Mitochondrial dysfunction and therapeutic approaches in respiratory and limb muscles of cancer cachectic mice. Exp Physiol. 2013;98(9):1349–65. Epub 2013/04/30. doi: 10.1113/expphysiol.2013.072496. PubMed PMID: 23625954.

38. VanderVeen BN, Hardee JP, Fix DK, Carson JA. Skeletal muscle function during the progression of cancer cachexia in the male Apc(Min/+) mouse. J Appl Physiol (1985). 2018;124(3):684–95. Epub 2017/11/11. doi: 10.1152/japplphysiol.00897.2017. PubMed PMID: 29122966; PMCID: PMC5899274.

39. Kang C, Ji LL. Muscle immobilization and remobilization downregulates PGC-1alpha signaling and the mitochondrial biogenesis pathway. J Appl Physiol (1985). 2013;115(11):1618–25. Epub 2013/08/24. doi: 10.1152/japplphysiol.01354.2012. PubMed PMID: 23970536.

40. Bowen TS, Rolim NP, Fischer T, Baekkerud FH, Medeiros A, Werner S, Bronstad E, Rognmo O, Mangner N, Linke A, Schuler G, Silva GJ, Wisloff U, Adams V, Optimex Study G. Heart failure with preserved ejection fraction induces molecular, mitochondrial, histological, and functional alterations in rat respiratory and limb skeletal muscle. Eur J Heart Fail. 2015;17(3):263–72. Epub 2015/02/07. doi: 10.1002/ejhf.239. PubMed PMID: 25655080.

41. Schmidt CA, Ryan TE, Lin CT, Inigo MMR, Green TD, Brault JJ, Spangenburg EE, McClung JM. Diminished force production and mitochondrial respiratory deficits are strain-dependent myopathies of subacute limb ischemia. J Vasc Surg. 2017;65(5):1504–14 e11. Epub 2016/12/28. doi: 10.1016/j.jvs.2016.04.041. PubMed PMID: 28024849; PMCID: PMC5403600.

42. van den Berg M, Hooijman PE, Beishuizen A, de Waard MC, Paul MA, Hartemink KJ, van Hees HWH, Lawlor MW, Brocca L, Bottinelli R, Pellegrino MA, Stienen GJM, Heunks LMA, Wust RCI, Ottenheijm CAC. Diaphragm Atrophy and Weakness in the Absence of Mitochondrial Dysfunction in the Critically Ill. Am J Respir Crit Care Med. 2017;196(12):1544–58. Epub 2017/08/09. doi: 10.1164/rccm.201703-0501OC. PubMed PMID: 28787181; PMCID: PMC5754442.

43. Ferreira R, Vitorino R, Alves RM, Appell HJ, Powers SK, Duarte JA, Amado F. Subsarcolemmal and intermyofibrillar mitochondria proteome differences disclose functional specializations in skeletal muscle. Proteomics. 2010;10(17):3142–54. Epub 2010/07/29. doi: 10.1002/pmic.201000173. PubMed PMID: 20665633.

44. Pham AH, McCaffery JM, Chan DC. Mouse lines with photo-activatable mitochondria to study mitochondrial dynamics. Genesis. 2012;50(11):833–43. Epub 2012/07/24. doi: 10.1002/dvg.22050. PubMed PMID: 22821887; PMCID: PMC3508687.

45. Duvall CL, Taylor WR, Weiss D, Guldberg RE. Quantitative microcomputed tomography analysis of collateral vessel development after ischemic injury. Am J Physiol Heart Circ Physiol. 2004;287(1):H302–10. Epub 2004/03/16. doi: 10.1152/ajpheart.00928.2003. PubMed PMID: 15016633.

46. Boerckel JD, Uhrig BA, Willett NJ, Huebsch N, Guldberg RE. Mechanical regulation of vascular growth and tissue regeneration in vivo. Proc Natl Acad Sci U S A. 2011;108(37):E674–80. Epub 2011/08/31. doi: 10.1073/pnas.1107019108. PubMed PMID: 21876139; PMCID: PMC3174614.

47. Akimoto T, Sorg BS, Yan Z. Real-time imaging of peroxisome proliferator-activated receptor-gamma coactivator-1alpha promoter activity in skeletal muscles of living mice. Am J Physiol Cell Physiol. 2004;287(3):C790–6. Epub 2004/05/21. doi: 10.1152/ajpcell.00425.2003. PubMed PMID: 15151904.

48. Nader GA, Esser KA. Intracellular signaling specificity in skeletal muscle in response to different modes of exercise. J Appl Physiol (1985). 2001;90(5):1936–42. Epub 2001/04/12. doi: 10.1152/jappl.2001.90.5.1936. PubMed PMID: 11299288.

49. Safdar A, Little JP, Stokl AJ, Hettinga BP, Akhtar M, Tarnopolsky MA. Exercise increases mitochondrial PGC-1alpha content and promotes nuclear-mitochondrial cross-talk to coordinate mitochondrial biogenesis. J Biol Chem. 2011;286(12):10605–17. Epub 2011/01/20. doi: 10.1074/jbc.M110.211466. PubMed PMID: 21245132; PMCID: PMC3060512.

50. Puigserver P, Wu Z, Park CW, Graves R, Wright M, Spiegelman BM. A cold-inducible coactivator of nuclear receptors linked to adaptive thermogenesis. Cell. 1998;92(6):829–39. Epub 1998/04/07. PubMed PMID: 9529258.

51. VanderVeen BN, Fix DK, Carson JA. Disrupted Skeletal Muscle Mitochondrial Dynamics, Mitophagy, and Biogenesis during Cancer Cachexia: A Role for Inflammation. Oxid Med Cell Longev. 2017;2017:3292087. Epub 2017/08/09. doi: 10.1155/2017/3292087. PubMed PMID: 28785374; PMCID: PMC5530417.

52. van Horssen J, van Schaik P, Witte M. Inflammation and mitochondrial dysfunction: A vicious circle in neurodegenerative disorders? Neurosci Lett. 2017. Epub 2017/07/03. doi: 10.1016/j.neulet.2017.06.050. PubMed PMID: 28668382.

53. Tidball JG. Mechanisms of muscle injury, repair, and regeneration. Compr Physiol. 2011;1(4):2029–62. Epub 2011/10/01. doi: 10.1002/cphy.c100092. PubMed PMID: 23733696.

54. Sadtler K, Estrellas K, Allen BW, Wolf MT, Fan H, Tam AJ, Patel CH, Luber BS, Wang H, Wagner KR, Powell JD, Housseau F, Pardoll DM, Elisseeff JH. Developing a pro-regenerative biomaterial scaffold microenvironment requires T helper 2 cells. Science. 2016;352(6283):366–70. Epub 2016/04/16. doi: 10.1126/science.aad9272. PubMed PMID: 27081073; PMCID: PMC4866509.

55. Tidball JG, Villalta SA. Regulatory interactions between muscle and the immune system during muscle regeneration. Am J Physiol Regul Integr Comp Physiol. 2010;298(5):R1173–87. Epub 2010/03/12. doi: 10.1152/ajpregu.00735.2009. PubMed PMID: 20219869; PMCID: PMC2867520.

56. Warren GL, Call JA, Farthing AK, Baadom-Piaro B. Minimal Evidence for a Secondary Loss of Strength After an Acute Muscle Injury: A Systematic Review and Meta-Analysis. Sports Med. 2017;47(1):41–59. Epub 2016/04/22. doi: 10.1007/s40279-016-0528-7. PubMed PMID: 27100114; PMCID: PMC5214801 assist in the preparation of this article. Conflicts of interest Gordon Warren, Jarrod Call, Amy Farthing, and Bemene Baadom-Piaro declare that they have no conflicts of interest relevant to the content of this review.

57. Wagatsuma A, Kotake N, Yamada S. Muscle regeneration occurs to coincide with mitochondrial biogenesis. Mol Cell Biochem. 2011;349(1-2):139–47. Epub 2010/11/27. doi: 10.1007/s11010-010-0668-2. PubMed PMID: 21110070.

58. Duguez S, Feasson L, Denis C, Freyssenet D. Mitochondrial biogenesis during skeletal muscle regeneration. Am J Physiol Endocrinol Metab. 2002;282(4):E802–9. Epub 2002/03/08. doi: 10.1152/ajpendo.00343.2001. PubMed PMID: 11882500.

59. Jash S, Adhya S. Induction of muscle regeneration by RNA-mediated mitochondrial restoration. FASEB J. 2012;26(10):4187–97. Epub 2012/07/04. doi: 10.1096/fj.11-203232. PubMed PMID: 22751011.

60. Nichenko AS, Southern WM, Atuan M, Luan J, Peissig KB, Foltz SJ, Beedle AM, Warren GL, Call JA. Mitochondrial maintenance via autophagy contributes to functional skeletal muscle regeneration and remodeling. Am J Physiol Cell Physiol. 2016;311(2):C190–200. Epub 2016/06/10. doi: 10.1152/ajpcell.00066.2016. PubMed PMID: 27281480.

61. Beltran MJ, Burns TC, Eckel TT, Potter BK, Wenke JC, Hsu JR, Skeletal Trauma Research C. Fate of combat nerve injury. J Orthop Trauma. 2012;26(11):e198–203. Epub 2012/03/23. doi: 10.1097/BOT.0b013e31823f000e. PubMed PMID: 22437422.

62. Horn A, Van der Meulen JH, Defour A, Hogarth M, Sreetama SC, Reed A, Scheffer L, Chandel NS, Jaiswal JK. Mitochondrial redox signaling enables repair of injured skeletal muscle cells. Sci Signal. 2017;10(495). Epub 2017/09/07. doi: 10.1126/scisignal.aaj1978. PubMed PMID: 28874604.

63. Vila MC, Rayavarapu S, Hogarth MW, Van der Meulen JH, Horn A, Defour A, Takeda S, Brown KJ, Hathout Y, Nagaraju K, Jaiswal JK. Mitochondria mediate cell membrane repair and contribute to Duchenne muscular dystrophy. Cell Death Differ. 2017;24(2):330–42. Epub 2016/11/12. doi: 10.1038/cdd.2016.127. PubMed PMID: 27834955; PMCID: PMC5299714.

64. Sandri M, Lin J, Handschin C, Yang W, Arany ZP, Lecker SH, Goldberg AL, Spiegelman BM. PGC-1alpha protects skeletal muscle from atrophy by suppressing FoxO3 action and atrophy-specific gene transcription. Proc Natl Acad Sci U S A. 2006;103(44):16260–5. Epub 2006/10/21. doi: 10.1073/pnas.0607795103. PubMed PMID: 17053067; PMCID: PMC1637570.

65. Andersen CL, Jensen JL, Orntoft TF. Normalization of real-time quantitative reverse transcription-PCR data: a model-based variance estimation approach to identify genes suited for normalization, applied to bladder and colon cancer data sets. Cancer Res. 2004;64(15):5245–50. Epub 2004/08/04. doi: 10.1158/0008-5472.CAN-04-0496. PubMed PMID: 15289330.

66. Kuznetsov AV, Veksler V, Gellerich FN, Saks V, Margreiter R, Kunz WS. Analysis of mitochondrial function in situ in permeabilized muscle fibers, tissues and cells. Nat Protoc. 2008;3(6):965–76. Epub 2008/06/10. doi: 10.1038/nprot.2008.61. PubMed PMID: 18536644.

67. Srere PA. Citrate Synthase. Methods Enzymol: Academic Press; 1969. p. 3–11.

68. Little RV, Brubaker RR. Characterization of deoxyribonucleic acid from Yersinia pestis by ethidium bromide-cesium chloride density gradient centrifugation. Infect Immun. 1972;5(4):630–1. Epub 1972/04/01. PubMed PMID: 4636787; PMCID: PMC422416.

69. Aihara H, Miyazaki J. Gene transfer into muscle by electroporation in vivo. Nat Biotechnol. 1998;16(9):867–70. Epub 1998/09/22. doi: 10.1038/nbt0998-867. PubMed PMID: 9743122.

70. Tehrani KF, Zhang Y, Shen P, Kner P. Adaptive optics stochastic optical reconstruction microscopy (AO-STORM) by particle swarm optimization. Biomed Opt Express. 2017;8(11):5087–97. Epub 2017/12/01. doi: 10.1364/BOE.8.005087. PubMed PMID: 29188105; PMCID: PMC5695955.

